# Barley Can Utilise Algal Fertiliser to Maintain Yield and Malt Quality Compared to Mineral Fertiliser

**DOI:** 10.1101/2025.08.12.667670

**Authors:** David James Ashworth, Stefan Masson, Tom Mulholland, Davide Bulgarelli, Kelly Houston

**Author notes:** **Correspondence:** Dr Kelly Houston.

## Abstract

**Background and Aims:** Mineral fertilisers are essential for global crop productivity. However, their use is unsustainable due to high energy costs from sourcing and processing them, and nitrous oxide emissions after their application. An alternative proposition is the use of microalgal biomass as a nutrient source for fertilisation. To date, there is limited data on how the productivity of barley grown with microalgae would compare to barley grown with mineral fertilisers. Furthermore, the impact of the fertiliser source on suitability of grain for malting, one of barley’s main end uses, should be considered.

**Methods:** We quantified the phenology, reflectance indices and yield components (including yield itself) of barley cv. Laureate grown with either *Chlorella vulgaris* powder or mineral fertiliser in three glasshouse trials in different soils, to assess the robustness of our findings under soil-to-soil variation. We extended this approach to carry out a field trial from which we malted the grain collected at the end of the season and quantified malt quality.

**Results:** We hypothesised that there would be no statistically significant difference in plant performance when grown with microalgae or mineral fertiliser when nitrogen addition was equal. Both glasshouse trials and a field trial showed that plants grown with microalgae and mineral fertiliser do not significantly differ in any yield component, yield itself, or measure of malt quality except total malt nitrogen.

**Conclusions:** Supplying the same amount of nitrogen to plants via microalgal biomass instead of mineral fertiliser produced barley with the same yield and malt quality as plants grown with mineral fertiliser.

## 1 Introduction

Mineral fertiliser has become ubiquitous in non-organic farming, due to its ability to increase yield by an estimated 50% (International Fertilizer Association 2018). However, the macronutrients found within mineral fertiliser, nitrogen (N), phosphorus (P) and potassium (K), have problems stemming from both sourcing and application. The Haber-Bosch Process, responsible for nitrogenous fertiliser, is estimated to consume between 1-3% of total global energy and contribute 1-3% of total greenhouse gas emissions (Institute for Industrial Productivity 2013). Phosphorus and K also contribute to ecological and climatic problems, as they are mined as rock phosphate and potash, respectively (Al Rawashdeh 2020; Walan *et al*. 2014). This both damages the environment through mining and generates large volumes of CO_2_ through energy and fuel consumption for extraction and processing. In addition to their production, the application of mineral fertilisers has further drawbacks. Applying nitrogenous mineral fertiliser further contributes to climate change as it decomposes to N_2_O, which accounts for 4-5% of global total greenhouse gas emissions (International Fertilizer Association 2018). Furthermore, N and P applied in excess of what plants can take up may be dissolved in rainwater or water provided by irrigation and subsequently be washed off into local watercourses. Increased nutrients in the watercourses can then cause eutrophication, which is harmful to the local ecosystems and to humans should drinking water be contaminated or sources of food be affected (Dodds *et al*. 2016; Hwang 2020; Le Moal *et al*. 2019).

Mineral fertiliser must be replaced, or its use greatly reduced, if humanity is to mitigate its detrimental impacts on the planet while maintaining or increasing crop yields. One proposed replacement for mineral fertiliser are products derived from eukaryotic microalgae or cyanobacteria that are safe for use in agriculture and do not accumulate toxins (hereafter collectively referred to as microalgae). Two such classes of microalgae derived product are microbial biofertilisers (MBFs), which are living microalgae (or other microbes) that can improve the availability of macro- and micronutrients in soil, and microbial biostimulants (MBSs), which are extracts or whole microalgae that can increase plant nutrient uptake, growth or quality through non-nutrient compounds affecting the plant’s physiology (Abdel-Raouf 2012; du Jardin 2015; Kapoore *et al*. 2021; Kawalekar 2013; Ronga *et al*. 2019). These two classes of microalgae derived products ignore a third class of product, those which use microalgae, living or dead at the time of application, to directly provide the macro- and micronutrients necessary for plant growth *from their own biomass*. These have no formal name, though are commonly put under the blanket term ‘organic fertiliser’ or ‘algal nutrients’ in papers that test these products (Alobwede *et al*. 2019; Mau *et al*. 2022; Schreiber *et al*. 2018). These products are the focus of this study, and we will refer to them as microalgal fertilisers. Microalgal fertilisers, MBFs and MBSs have all been proven to be effective at sustaining or increasing one or more aspects of plant health or agronomy (Kapoore *et al*. 2021; Ronga *et al*. 2019).

There are advantages to using microalgal fertilisers beyond sustaining plant performance. Starting with their production, they do not necessarily rely on the Haber process or mining to provide their N, P and K; instead, nutrient rich wastewaters can be used as growth media.

Multiple different species of microalgae, both prokaryotic, such as *Spirulina/Arthrospira platensis* G. (Lima e Silva *et al*. 2024; Wuang *et al*. 2016) and eukaryotic, such as *Chlorella* sp. B. or *Chlamydomonas reinhardtii* D., have been grown on different sources of wastewater (Mohsenpour *et al*. 2021; Nguyen *et al*. 2022). Among the microalgae tested for their ability to treat wastewater, and the genus of importance to this study, species of *Chlorella* have been grown on a range of waste waters including, but not limited to, synthetic wastewater as a proof of concept (Zhao *et al*. 2018), municipal wastewater (Amaya-Santos *et al*. 2022; Mennaa *et al*. 2015; Morillas-Espana *et al*. 2022; Ranglová *et al*. 2021; Wang *et al*. 2010; Zhou *et al*. 2018), animal slurry and its anaerobic digestate (Zhou *et al*. 2018), waste aquaculture water and industrial food processing waste (Mukherjee *et al*. 2015; Viegas *et al*. 2021; Zhou *et al*. 2018) due to their very high N (>90% in most studies), P (>80% in most studies) and micronutrient removal efficiencies (Mennaa *et al*. 2015; Morillas-Espana *et al*. 2022; Mukherjee *et al*. 2015; Viegas *et al*. 2021; Wang *et al*. 2010; Zhao *et al*. 2018; Zhou *et al*. 2018). In addition to efficient recovery of macro- and micronutrients, microalgal cultivation releases less greenhouse gas than traditional wastewater treatment (Nguyen *et al*. 2022). This is in partially because approximately 50% of the energy spent on wastewater treatment is oxygen delivery, which microalgae does not require (Mohsenpour *et al*. 2021). Furthermore, microalgae can absorb nitrogen compounds before they decompose to the greenhouse gas nitrous oxide, the generation of which is a concern in traditional wastewater treatment systems (Mohsenpour *et al*. 2021). However, it is also reported that microalgae can generate nitrous oxide (Burlacot *et al*. 2020; Plouviez *et al*. 2018) through the action of nitrate reductase producing NO, which is then reduced further to nitrous oxide (Bellido-Pedraza *et al*. 2022). Therefore, a detailed lifecycle analysis would be necessary to determine if there is a greenhouse gas saving for a particular implementation of a microalgal cultivation system. These energy and nitrous oxide emission savings, if present, could greatly reduce overall greenhouse gas emissions when using microalgae as a fertiliser if also linked to a reduction in the Haber process and P/K mining. When looking to application on farm, microalgal fertilisers are considered a source of slow-release N and P, with Coppens *et al*. (2015) reporting plant available N at 31% after 95 days (in line with commercial slow release fertilisers), plant available P at roughly 60% for the duration of the experiment, and phosphate P at 7% . Lower concentrations of soluble nutrients reduce runoff and so could result in less eutrophication. Additionally, there is evidence that use of microalgal fertilisers can significantly increase total soil carbon by up to 17% (Alobwede *et al*. 2019), meaning there is potential for both replenishment of degraded soils and nourishment of otherwise healthy soils (Abdel-Raouf 2012; Kawalekar 2013).

Cultivated barley (*Hordeum vulgare* L. ssp. *vulgare*) is a globally important cereal crop, being the 4^th^ most cultivated cereal behind maize, rice and wheat (FAOSTAT 2020). In the UK, the 9^th^ largest producer globally and 7^th^ in Europe (FAOSTAT 2020), barley is the second most produced crop by mass, behind wheat and ahead of rapeseed, contributing £1.16 billion to the economy (DEFRA 2021). Barley grown in the UK has two main markets. The first is the premium market of brewing and distilling, which uses roughly 25% of the grain, and adds £5.5 billion to the economy from Scotch Whisky (DEFRA 2021; Scotch Whisky Association 2021). The remaining 75% of barley goes almost entirely to animal feed. The use of barley in brewing and distilling is of particular interest to this study, as both processes generate a large volume of wastewater, which must be treated by both UK and Scottish law if it is to be released into sewage or watercourses (Scottish Government 2012). General distillery and brewery wastewater is currently treated by a range of processes including physical, chemical, physicochemical, and biological (Amenorfenyo *et al*. 2020; Ghosh Ray *et al*. 2018; Kharayat 2012; Laurinavichene *et al*. 2018; Pant *et al*. 2007; Ratna *et al*. 2021; Sonawane *et al*. 2014; Yamasaki *et al*. 2006). The biological treatment methods include anaerobic digestion (which can generate bio-methane), aerobic digestion, algal growth, or plant growth through simulated wetlands. Species of *Chlorella* have been shown to be suitable for use in biological treatment of waste from a sugar mill, which produces ethanol (Valderrama *et al*. 2002), and cassava ethanol waste (Yang *et al*. 2008) in both axenic and non-axenic culture. The ability of microalgae to treat distillery wastewater alone or in consortia, such as with *Lemna minuscula* (Valderrama *et al*. 2002) is evidence of their value as wastewater remediators, and another reason why uses for this microalgal biomass, such as for use as fertiliser, should be investigated.

This study seeks to investigate the suitability of microalgae as a fertiliser for barley, and the impact of this type of fertiliser on barley malt quality. Establishing a microalgal cultivation system capable of producing the masses needed for these experiments is a significant time, money and energy commitment. Therefore, to facilitate proof of concept experiments we used purchased microalgae as a fertiliser instead of growing our own, with the view to use microalgae grown on distillery waste in the future. We compared plants grown using microalgal fertiliser with a negative control, where we did not provide any additional inputs to the soil (no additions) and a mineral fertiliser positive control. Our comparisons were based on the phenology, yield components, grain quality, and malt quality of spring barley cv. Laureate grown with these different treatments. We grew barley in both glasshouse trials, to allow assessment of the performance of microalgal fertilisers in controlled conditions, and field trials, to determine efficacy of microalgal fertilisers in a scenario closer to the intended end user, extending the work done previously on wheat and barley that had not reached maturity (Lima e Silva *et al*. 2024; Schreiber *et al*. 2018). We hypothesised that, when the amount of nitrogen provided by each fertiliser treatment is equal and at the recommended rate for malting barley, 118kg(N)/ha (Kendall *et al*. 2021), there will be no significant differences in yield components, grain quality or malt quality between plants grown with microalgae or mineral fertiliser, showing that microalgae can be as effective as mineral fertiliser for producing barley for malting.

## 2 Materials and Methods

### 2.1-3 Glasshouse experiments

### 2.1 1^st^ Algae Response Experiment

#### 2.1.1 Plant Growth Conditions

Four treatments were prepared using Bullionfield soil taken from a field (56.460253, - 3.071053) local to the James Hutton Institute, Dundee. All soils used for these experiments were analysed by Yara UK using their SA10 (soil mineral nitrogen) and BSE SOL (broad spectrum soil health) testing suites (Online Resource 1, **Table S1**). Bullionfield soil was categorised by Yara UK as a sandy silt loam, with a pH of 5.8 and a C:N of 12.4. After excavation and before any further processing, soil was passed through a 10mm sieve, to remove large stones which would occupy a disproportionate volume when potted and may affect drainage. The sieved soil was then given one of four treatments. The first treatment was no additional nutrients supplied to the soil to function as a negative control, designated No Additions. The second treatment was the addition of dried *Chlorella vulgaris* B. (hereafter referred to a *C. vulgaris*) powder that provided an addition of 94.3 kg(N)/ha, designated

Algae Only. 94.3kg(N)/ha was chosen as the dose, as based on analysis of soil from a field near the sampling site for Bullionfield, it brought the total N (soil mineral N plus applied N) up to 118kg(N)/ha, which is AHDB’s recommended rate for malting barley (Kendall *et al*. 2021). As the *C. vulgaris* provided less K and Ca than the positive control when normalising for N, the third treatment was *C. vulgaris* powder that provided an addition of 94.3 kg(N)/ha with additional K (as KCl) and Ca (as CaCl_2_.2H_2_O), designated Algae Supplement.

Potassium and Ca were added to bring the concentration of these nutrients up to the level seen in the final treatment, which was addition of mineral salts to the soil providing 94.3 kg(N)/ha, following the molar ratios found in a modified Hoagland solution recipe (Hoagland 1950) as a positive control, designated Mineral, with the exclusion of the micronutrients, as such small masses of salt would not homogenously disperse in the soil (Online Resource 1, **Table S2**). We chose to use the salts and their molar ratios provided by the Hoagland solution recipe as the nutrients found in Hoagland solution are well known to support plant growth.

Note, we did not apply Hoagland solution itself, only used the recipe to provide the salts and their molar ratios. Treatments are summarised in Online Resource 1, **Fig. S1**.

*C. vulgaris* was chosen as the microalgal fertiliser for these experiments as it has a long history of safe use in food and nutraceuticals. Furthermore, *C. vulgaris* is not classed as a ‘novel food’ under Regulation (EU) 2015/2283 (Czech Republic 2022), meaning there is no regulatory barrier to use on-farm. Additionally, it accumulates high concentrations of N, P and other macro and micronutrients (Alobwede *et al*. 2019), and has a wide body of literature proving its efficacy in other plants (see Introduction). For this experiment, we used *C. vulgaris* batch one, which was purchased from Raw Living (Raw Living Ltd, UK), a nutraceutical company. The elemental compositions of all batches of *C. vulgaris* used in this study were analysed by tin catalysed combustion, with helium as the carrier gas, in a Thermo Finnigan FlashEA 1112 Elemental Analyzer (Thermo Fischer Scientific Inc, USA) for C and N, and nitric acid digestion followed by ICP-OES on an Avio 500 ICP-OES (Perkin Elmer, USA) for all other listed elements (Online Resource 1, **Table S3** and **Fig. S2**). All Raw Living sourced *C. vulgaris* was cultivated by the company in raceway ponds with a ‘food grade’ mineral nutrient solution as growth media prior to harvesting, drying and shipping (Raw Living, Personal Correspondence).

Each treatment was assigned to 12 “6 inch rose pots” (top diameter 15cm, depth 20cm) with each pot containing 2.9kg of dry soil. One seven-day old seedling of barley cv. Laureate, a two-row spring variety that is the current dominant malting variety in Scotland (AHDB Data and Analysis Team 2024), was planted into each pot. Two pots of each treatment were paired to give six pairs per treatment. Each pair was laid out in a complete block design with six blocks, one pair per treatment per block. Pots were individually placed in plastic dishes to ensure any excess water would pool and be reabsorbed and no nutrients would be lost. The seeds were germinated 03/02/2023 and were transferred to soil 10/02/2023. Plants were grown in a glasshouse at the James Hutton Institute, Dundee, set to 18⁰C day, 14⁰C night, 16h light, 8h dark with supplemental lighting as necessary to maintain a minimum of 200 µmol photons m^−2^ s^−1^ during the day period.

Each plant pot was watered to 80% field capacity with tap water three times a week for the first three weeks. From the fourth week onward, plants were watered to 80% field capacity twice a week, with one day a week being replaced with 72mL of 1x Micronutrient solution (Online Resource 1, **Table S2**), as the mineral treatment provided no additional trace micronutrients as solid salts, plus 178mL tap water for mineral treatment plants, or 250mL tap water for no additions, Algae Only or Algae Supplement plants. A picture of the experiment in early growth can be found in the supplementary material (Online Resource 1, **Picture S1**).

#### 2.1.2 Measurements of Development and Morphology

Three times a week, plant development was scored using the Zadoks Scale (Zadoks *et al*. 1974), and the number of tillers were counted. At 38 days after sowing (DAS), when plants were at GS22-25 (tillering), one plant per pair was harvested. The transmission spectrum from the middle of the 3^rd^ leaf on the main tiller was recorded, using the PolyPen RP410 UVIS (Photon Systems Instruments, Czechia), after which we removed the 3^rd^ leaf from the plant. The transmission spectrum was used to calculate the Normalised Difference Vegetation Index (NDVI) (Rouse *et al*. 1974). We measured the length of the 3^rd^ leaf by hand, the area of the 3^rd^ leaf using ImageJ (Schneider *et al*. 2012), and collected the aboveground biomass from each plant then dried it at 60⁰C until no further decrease in mass was seen, after which it was weighed to determine aboveground dry biomass.

The aboveground biomass was harvested from the remaining plant of each pair at maturity. Ears and vegetative tissues were separated, dried at 60⁰C and weighed to determine the aboveground dry biomass. Ear length was measured using ImageJ, and the number of filled grains counted. Ears were then threshed and, after removing any broken grain, we passed grain from each plant through a MARViN ProLine (MARViTECH GmbH, Germany) to determine mean grain length, width and area, and thousand grain weight (TGW).

### 2.2 2^nd^ Algae Response Experiment

This experiment was designed and carried out as described in section 2.1, with the following exceptions. A different soil designated Quarryfield 4 (56.453737, -3.076683, Online Resource 1, **Table S1**) from near the James Hutton Institute, Dundee, was used. Quarryfield 4 was categorised as a sandy loam with a pH of 6.2. and a C:N of 14. The algae and mineral treatments provided an additional 82.6 kg(N)/ha to reach a target soil nitrogen of 118 kg(N)/ha, and the algae supplement treatment had additional K, Ca, Mg, S and Fe to match the concentrations found in the mineral treatment, in the form of KCl, CaCl_2_.2H_2_O, CaSO_4_.2H_2_O, MgSO_4_ and FeNaEDTA, with the sulphate and chloride salts of Ca used to ensure the molar ratios of Ca and S were unchanged with regard to the other nutrients. Each treatment was assigned to 16 plants (eight pairs of pots). Seeds were germinated on 02/05/2023 and were transferred to soil on 10/05/2023. When over half of the plants were at GS22/had two tillers, we harvested one member of each pair. This was in place of harvesting at a fixed day after sowing, to account for differences in speed of development between experiments.

### 2.3 3^rd^ Algae Response Experiment

This experiment was designed as in section 2.2 with the following exceptions. A different batch of soil, designated Quarryfield 5, taken from the same field as Quarryfield 4, near the James Hutton Institute, Dundee, was used (Online Resource 1, **Table S1**). Quarryfield 5 was categorised as a sandy loam with a pH of 5.8 and a C:N of 11.9. The algae and mineral treatments provided an additional 38.96kg(N)/ha, to make up the 79.04kg(N)/ha already in the soil to 118kg(N)/ha. Batch two of algae powder, purchased from Raw Living, was used for this experiment (Online Resource 1, **Table S3, Fig. S2**). 16 seedlings per treatment were planted and grouped into eight pairs. Seeds were germinated on 11/07/2023 and were transferred to soil on 18/07/2023.

### 2.4 Spring 2024 Field Trial

#### 2.4.1 Planting

We performed a field trial comparing the phenology and yield components of cv. Laureate when grown with no additional nutrients, *C. vulgaris* pellets (algae pellets) or 24-4-14 NPK granules. This experiment was sown at Balruddery Farm, Angus, located close to the James Hutton Institute, Dundee (Hutchens field, 56.485530, -3.110772, Online Resource 1, **Table S1**), in a sandy silt loam following vining peas. The Hutchens soil had a pH of 6.3 and the C:N was 10. We filled 27 packets (one packet per experimental plot) with 185g of cv.

Laureate, to achieve a seed rate of 360 seeds/m^2^. Nine packets received no additions to the seed. Nine packets additionally received 224g NPK granules (Online Resource 1, **Table S4**, **Fig. S3**). The remaining nine packets were paired with separate packets containing 563g of *C. vulgaris* pellets (Online Resource 1, **Table S4**, **Figure S3**), as the pellets and seeds would not all fit in one packet. Pellets were chosen instead of powder for ease of handling by farm machinery. The algae pellets and NPK granules both provided an additional 61.6kg(N)/ha to bring the 56.4kg(N)/ha in the top 90cm of soil up to 118kg(N)/ha. The no additions and NPK plots were sown in one pass, while the plots receiving algae had seed sown first, then had a second pass to sow the algae pellets. Plots were 6m long and 1.5m wide with seeds and fertiliser sown in 8 parallel lines per plot. Treatments were assigned in three blocks, each a 3×3 Latin square. The experimental plots were separated by guard plots of cv. LG Diablo, which received no additional fertiliser. The plots were sown on 26/04/2024. All plots received three standard fungicide and herbicide applications over the growing season. An aerial image of the trial can be found in the supplementary material (Online Resource 1, **Picture S2**).

#### 2.4.2 Development and Harvest Measurements

At 26DAS, we scored the plots for the emergence of plants. At 26, 35, 40 and 56DAS, the NDVI of each plot was recorded using the PolyPen RP410 UVIS (Photon Systems Instruments, Czechia). Ten plants per plot were chosen at random, their third leaf on the main tiller selected for measurement, and their values averaged to provide the mean plot NDVI. At 33, 40 and 54DAS, we recorded the median number of tillers and Zadoks score for each plot. We also determined flag leaf emergence date, when approximately half of the plot had a horizontal flag leaf, and heading date, when approximately half of the plot had visible ears.

At plant maturity but prior to harvest, we collected grab samples of the plants in the middle 20cm of the middle two rows of each plot. Grains per ear and ear length of three random ears per grab sample were recorded. We threshed the whole grab sample, dried the totality of the threshed grain (150-175 seeds per plot) at 60⁰C, then measured TGW and grain dimensions on the MARViN ProLine (MARViTECH GmbH, Germany). Grain from grab samples, as opposed to whole plots, were used for dimension measurement as it had not been passed over a sieve, giving a more accurate-to-field size distribution.

The field plots were harvested and threshed by combine harvester, which also measured mass of grain per plot. Grain from each plot was kept separate and dried briefly to remove surface moisture and prevent growth of mould. Approximately 1kg of grain from each plot was cleaned by passing the grain over a series of sieves, removing debris and leaving grain >2.5mm. The protein content and moisture of the cleaned sample was measured using an Infratec (Foss, Denmark). Protein content was converted to nitrogen by dividing by 5.75, the conversion factor recommended by AHDB (Sylvester-Bradley *et al*. 2009). Using the mass of grain per plot and the moisture content, we calculated yield of dry grain per plot.

#### 2.4.3 Malt Quality

Grain from the combine harvested (not grab) samples was malted, after it had been stored at 15⁰C for approximately five months. Each plot had 50g samples of grain malted in quadruplicate, to give four malting reps. Malting reps were distributed during malting as a random complete block design, where each tank acted as a block. Each sample was weighed into a water permeable tin, four of which could fit in one half of a drum, with four drums per malt tank, in two malt tanks running simultaneously (Curio Malting, formerly Custom Laboratory Products Ltd, UK). Samples were malted using the following regime, with all water and air rests at 15⁰C: 8hr steep, 10hr air rest, 6hr steep, 8hr air rest, 4hr steep, 2hr air rest. The steeping phase was followed by germination for 24hrs at 20⁰C, 24hrs at 18⁰C, 24hrs at 17⁰C and 24hrs at 16⁰C for a total malting time of 132hrs and 39minutes. Drums were rotated throughout the programme. The programme was run twice, each with two of the four malting reps. The germinated malt in each sample was kilned (Curio Malting, UK) immediately following the end of the germination phase, using the following regime: 6hrs at 50⁰C fan speed 70, 6hrs 55⁰C fan speed 70, 60⁰C until 7% air humidity fan speed 30, 65⁰C to 5% air humidity fan speed 30. After the malt was kilned, it was dressed (removing the shoots and roots by rubbing it in cloth) and passed over a 2.5mm sieve to remove the debris.

Malt was stored in sealed plastic bags at 15⁰C until analysis was carried out at Glen Keith Technical Centre (Chivas Brothers, UK). Each malting replicate was kept separate, and the entirety of each sample (∼46g) was ground to flour using a Bühler DLFU disc mill (Bühler, Switzerland) set to 0.2mm. 5.0000g ± 0.01g of flour was weighed into a metal dish, recording the exact mass. Each dish was heated in a Thermo Scientific Heratherm oven (Thermo Fischer Scientific Inc, USA) at 105⁰C for 3 hours, after which they were left to cool in a desiccator for 20 minutes, then weighed again to determine the moisture content of the malt. 25.00g of flour (not dried) from each sample was weighed into a mashing tin. Each tin had a 4.5cm magnetic stir bar added and was placed in a Rototherm CT4 Mashing Bath (Rototherm Canongate Technology, UK). Between 20 and 23 flour samples were mashed per run of the mashing bath, for a total of five mashes. Each tin was filled with 360 ± 10mL of 65⁰C demineralised water, then a watch glass placed on top to prevent evaporation. Flour samples were kept at this temperature and stirred for one hour, after which they were cooled to 20⁰C by replacement of the warm water surrounding the tins with cold water. The contents of each tin were added to separate brewer’s flasks (a 515mL volumetric flask) and the volume made up to 515mL with demineralised water. The contents of each brewer’s flask were filtered through Whatman V2 filter paper (Cytiva, USA) for 30 minutes. The resulting filtrate is the wort. Approximately 50mL of each wort was decanted into a vial and the specific gravity was measured on a DMA 5000M density meter (Anton Paar GmbH, Austria), which the associated software automatically converted to extract strength in Litre degrees/kg. During analysis we doubled this value, as the method as written and the specific gravity to extract strength calculation assumes 50g of flour. Doubling was validated as accurate by measuring the extract strength of the wort of 50g and 25g of the same test malt. 1mL of each wort, noting the exact mass, was decanted into a separate Elemental Microanalysis Evaporating Tin Foil Cup (Elemental Microanalysis Ltd, UK), and heated on a SB300 hot plate (Cadmus Products Ltd, UK) with the cup holding attachment at setting 3.5 until a dry, brittle crust formed. The nitrogen content of the wort (soluble nitrogen) was measured using a LECO FP828 (LECO Corporation, USA), corrected to 0% grain moisture then doubled to account for the 25g sample in a 50g method. Also using the LECO FP828, 1.4000-1.6000g of flour from each sample was weighed, noting the exact mass, into a LECO tin foil cup (LECO Corporation, USA), and the total nitrogen of the malt, corrected to 0% grain moisture, was measured. From the extract and total nitrogen, we calculated the predicted spirit yield (PSY) using Chivas Brothers’ proprietary equation for the 2024 harvest. Using the moisture content, we calculated the PSY on a dry basis.

### 2.5 Statistical Analysis

For all analyses and models fitted, diagnostic plots were examined for homoscedasticity and normality of residuals. If satisfactory, the appropriate parametric tests were performed, either ANOVA followed by Tukey’s HSD for cases where only the treatment was significant, or mixed model ANOVA followed by mixed model contrasts where a factor other than the treatment was significant. All analyses of data collected from separate experiments used linear mixed models in which treatment was modelled as a fixed effect and block nested within experiment as a random effect. Use of mixed models with experiment as a random factor was validated by performing standard ANOVAs for treatment, experiment and their interaction term. For all measurements, the interaction terms between treatment and experiment were not statistically significant (p > 0.05), implying the treatments had the same effect for each experiment, and therefore the treatment effect across experiments could be compared using mixed models.

If residual variance was found to increase with increasing fitted values, a log transformation was applied to the response variable, diagnostic plots re-examined, and the appropriate parametric tests conducted. Where a log transformation has been applied, a log axis is also provided on the graph and the p-value reported is from the log transformed model. If the response could not be transformed to give residuals suitable for parametric testing, the Kruskal-Wallis test followed by Dunn test with Benjamini-Hochberg correction for multiple testing was performed.

All analysis and graphing was conducted using R version 4.5.3 and R studio version 2026.01.2+418 (Posit Team 2024; R Core Team 2026). We used the packages “car” (Fox *et al*. 2019), “FSA” (Ogle *et al*. 2025), “ggplot2” (Wickham 2016), “ggpubr” (Kassambara 2023), “ggbeeswarm” (Clarke *et al*. 2023), “ggthemes” (Arnold 2024), “lme4” (Bates *et al*. 2015), “afex” (Singmann *et al*. 2024), “fmsb” (Nakazawa 2024), “tidyr” (Wickham *et al*.

2024), “stringr” (Wickham 2023), “extrafonts” (Chang 2023), “patchwork” (Pendersen 2025) and all their dependencies. All scripts can be found at https://github.com/DJAshworth1015/Barley-Can-Utilise-Algal-Feriliser-To-Maintain-Yield-And-Malt-Quality.

## 3 Results

### 3.1 Responses to Algal Fertiliser in Glasshouse Trials

#### 3.1.1 Phenology

In the first glasshouse experiment, there was a significant overall effect of treatment at 26 Days After Sowing (DAS, Kruskal-Wallis, p = 0.011). Plants grown with the Algae Only treatment produced tillers significantly faster than those with the Algae Supplement treatment (**Table 1**, Online Resource 1, **Fig. S4 (a)**). 38DAS also showed a significant effect of treatment (Kruskal-Wallis, p = 0.020). Plants grown with the Algae Only treatment produced tillers significantly faster than those grown with no additions at 38DAS (**Table 1**, Online Resource 1, **Fig. S4 (b)**). The scoring points either side of the reported days showed no significant differences in Zadoks score between treatments.

**Table 1.** Significance of comparisons for the glasshouse experiments in Online Resource 1. Graphs for each trait listed can be found in Online Resource 1, with figure numbers listed in Sections 3.1.1 and 3.1.2. For methods of obtaining p-values, KW = Kruskal-Wallis followed by Dunn Test, MMA = Mixed model ANOVA followed by mixed model contrasts. Different letters represent significance groups at p < 0.05, with multiple letters representing non-significant intermediates (*i.e.* ab is not significantly different from a or b, which are significantly different from one another). The “^” sign represents the highest value in the comparison (*e.g.* Algae Only has a higher Zadoks Score (GS) than Algae Supplement at 26DAS). Measurements were taken at GS22 unless otherwise stated

| Trait | P-value | No Additions | Algae Only | Algae Supplement | Mineral |
| --- | --- | --- | --- | --- | --- |
| 26DAS GS | KW 0.011 | ab | a<sup>^</sup> | b | ab |
| 38DAS GS | KW 0.02 | a | b<sup>^</sup> | ab | ab |
| 42DAS GS | KW 0.027 | a | ab | b<sup>^</sup> | ab |
| 45DAS GS | KW 0.048 | a | ab | b<sup>^</sup> | ab |
| 46DAS GS | KW 0.007 | a | ab | b<sup>^</sup> | a |
| 66DAS GS | KW 0.029 | a | ab | b<sup>^</sup> | ab |
| 3<sup>rd</sup> leaf length | MMA 0.528 | a | a | a | a |
| 3<sup>rd</sup> leaf area | MMA 0.870 | a | a | a | a |
| Aboveground dry biomass | MMA 0.708 | a | a | a | a |
| NDVI | MMA 0.345 | a | a | a | a |

At 42DAS (Kruskal-Wallis for treatment, p = 0.027), plants grown with the Algae Supplement treatment entered stem elongation (node formation) significantly earlier than plants grown with no additions. The significant effect of treatment was maintained across 45DAS (Kruskal-Wallis, p = 0.048) and 46DAS (Kruskal-Wallis, p = 0.0070), where the plants grown with Algae Supplement had significantly more nodes than those grown with no additions, and the Algae Supplement treated plants also developed nodes significantly faster than the mineral treatment at 46DAS (**Table 1**, Online Resource 1, **Fig. S5 (a), (b),** and **(c)**). The difference between the Algae Supplement treated plants and the other treatments became non-significant at the next timepoint (48DAS, data not shown). The only other timepoint at which there was a significant difference between treatments was at 66DAS (Kruskal-Wallis, p = 0.029), where the ears of the Algae Supplement treated plants started to emerge from the flag leaf sheath significantly faster than those of the plants grown with no additions (**Table 1**, Online Resource 1, **Fig. S5 (d)**).

In both the second and third glasshouse experiments, there were no significant differences in Zadoks score between treatments at any point during development.

#### 3.1.2 Morphometrics and Reflectance in Early Tillering

There were no significant differences between plants of cv. Laureate grown with no additions, Algae Only, Algae Supplement, or mineral salts in the third leaf length, third leaf area or aboveground dry biomass at GS21-23/early tillering (**Table 1**, Online Resource 1, **Fig. S6 (a), (b)** and **(c)**). Likewise, there are no significant differences in NDVI between any of the treatments at this sampling point (**Table 1**, Online Resource 1, **Fig. S6 (d)**). The lack of a significant difference between the negative control (no additions) and the positive control (mineral fertiliser) suggests that at this point in development the available nutrients are not a limiting factor for growth, nor is the ability to absorb photosynthetically active red light.

#### 3.1.3 Morphometrics and Grain Traits at Maturity

Overall, when combining the morphometric and grain trait data that we collected, we observed that plants grown using microalgal fertiliser are not significantly different from those grown with mineral fertiliser in every measured trait. Furthermore, plants grown using microalgal fertiliser have significantly greater earless dry biomass, number of tillers, filled grain, TGW, grain width and grain area than those grown with no additions.

At maturity, plants grown with the Algae Only, Algae Supplement, or mineral treatment did not have a significantly different earless dry biomass from one another, and plants grown in any of these three treatments had a significantly higher earless dry biomass than plants grown with no additions (Mixed model ANOVA, p < 0.001, **Fig. 1 (a)**). Plants grown with either microalgae or mineral fertiliser did not have a significantly different number of tillers from each other, and the number of tillers for plants grown with any of the three treatments significantly exceeded those grown with no additions (Mixed model ANOVA, p < 0.001, **Fig. 1 (b)**). Total filled grain per plant followed the same pattern as earless dry biomass, with the two microalgae treatments and the mineral treatment not significantly differing from one another, and plants grown with any of the three treatments filling significantly more grain than plants grown with no additions (Mixed model ANOVA, p = 0.007, **Fig. 1 (c)**). The number of filled grains per ear did not differ significantly between treatments (Mixed model ANOVA, p = 0.969, **Fig. 1 (d)**), indicating that the increases in the total number of filled grain in the microalgae and mineral treatments were driven by an increased number of tillers only.

**Fig. 1.**
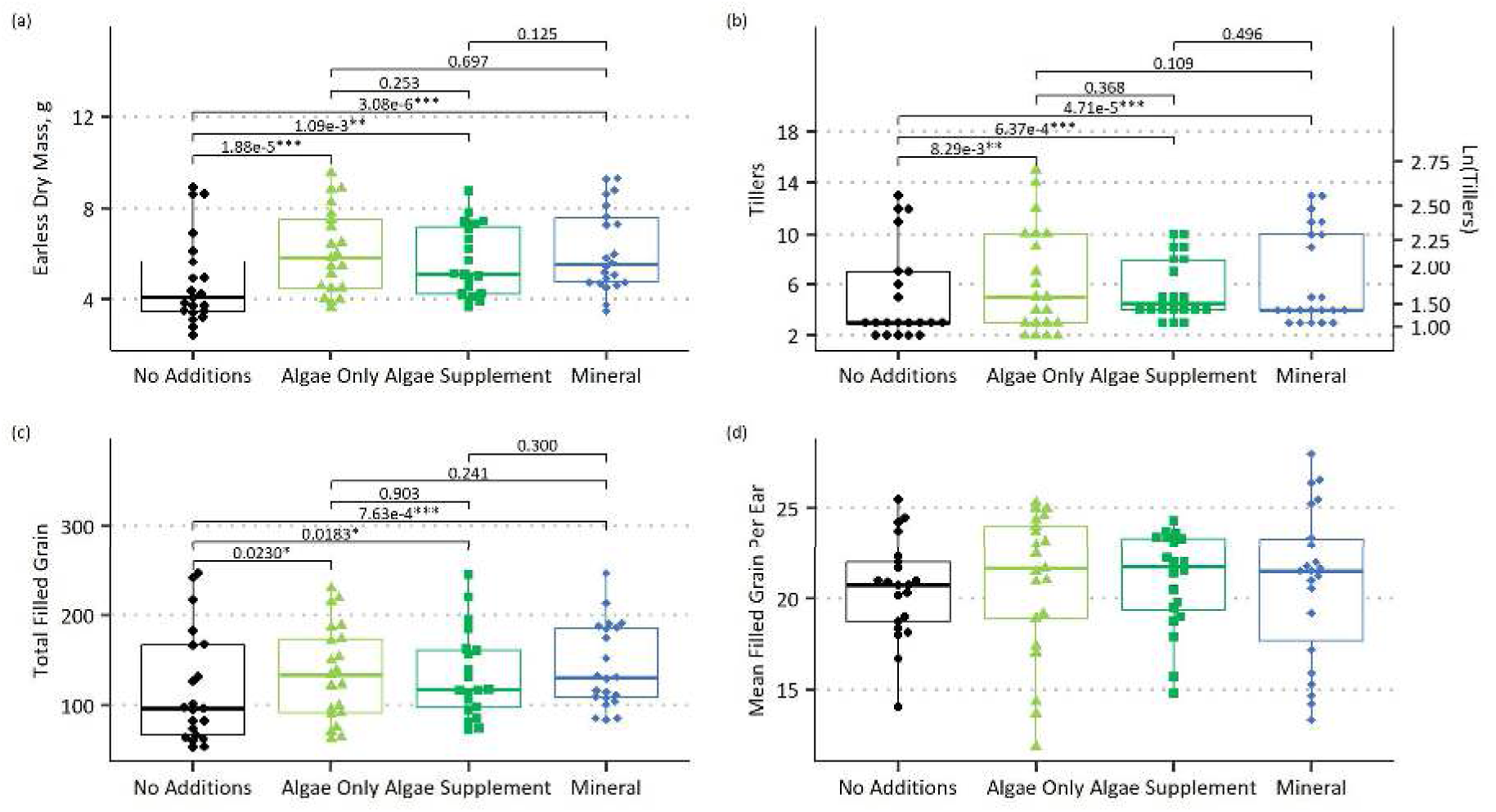
Morphometrics of cv. Laureate at maturity when grown with four different nutrient sources across three glasshouse experiments. **(a)** Earless dry mass. **(b)** Number of tillers. **(c)** Total filled grain per plant. **(d)** Mean filled grain per ear per plant. For all boxplots, the thick horizontal line is the median, thin horizontal lines are the 1st and 3rd quartile. Whiskers extend to the range, excluding outliers (>1.5x the interquartile range below quartile 1 or above quartile 3). Overall significance determined by mixed model ANOVA, p-value reported in the main text. Pairwise significance determined by mixed model contrasts. For **(b)**, the model fitted used the natural log of the tiller number, but untransformed tiller number is presented here for ease of interpretation. N = 21 for no additions, 21 for algae only, 20 for algae supplement, 22 for mineral. Each rep is one data point

When analysing the grain morphology traits, there were no significant differences between plants grown with either microalgae treatment and mineral fertiliser in thousand grain weight (TGW), and all three treatments produced plants with significantly greater TGWs than plants grown with no additions (Mixed model ANOVA, p = 0.008, **Fig. 2 (a)**). Grain from plants grown with the Algae Only treatment or mineral fertiliser did not differ significantly in width and had significantly wider grain than plants grown with no additions, with the Algae Supplement treatment being a non-significant intermediate (Mixed model ANOVA, p = 0.029, **Fig. 2 (b)**). There were no significant differences in grain length between any of the treatments (Mixed model ANOVA, p = 0.197, **Fig. 2 (c)**). Grain area was significantly larger for plants grown with the Algae Only treatment relative to those grown with the no additions treatment, with the Algae Supplement and mineral treatment being non-significant intermediates (Mixed model ANOVA, p = 0.018, **Fig. 2 (d)**).

**Fig. 2.**
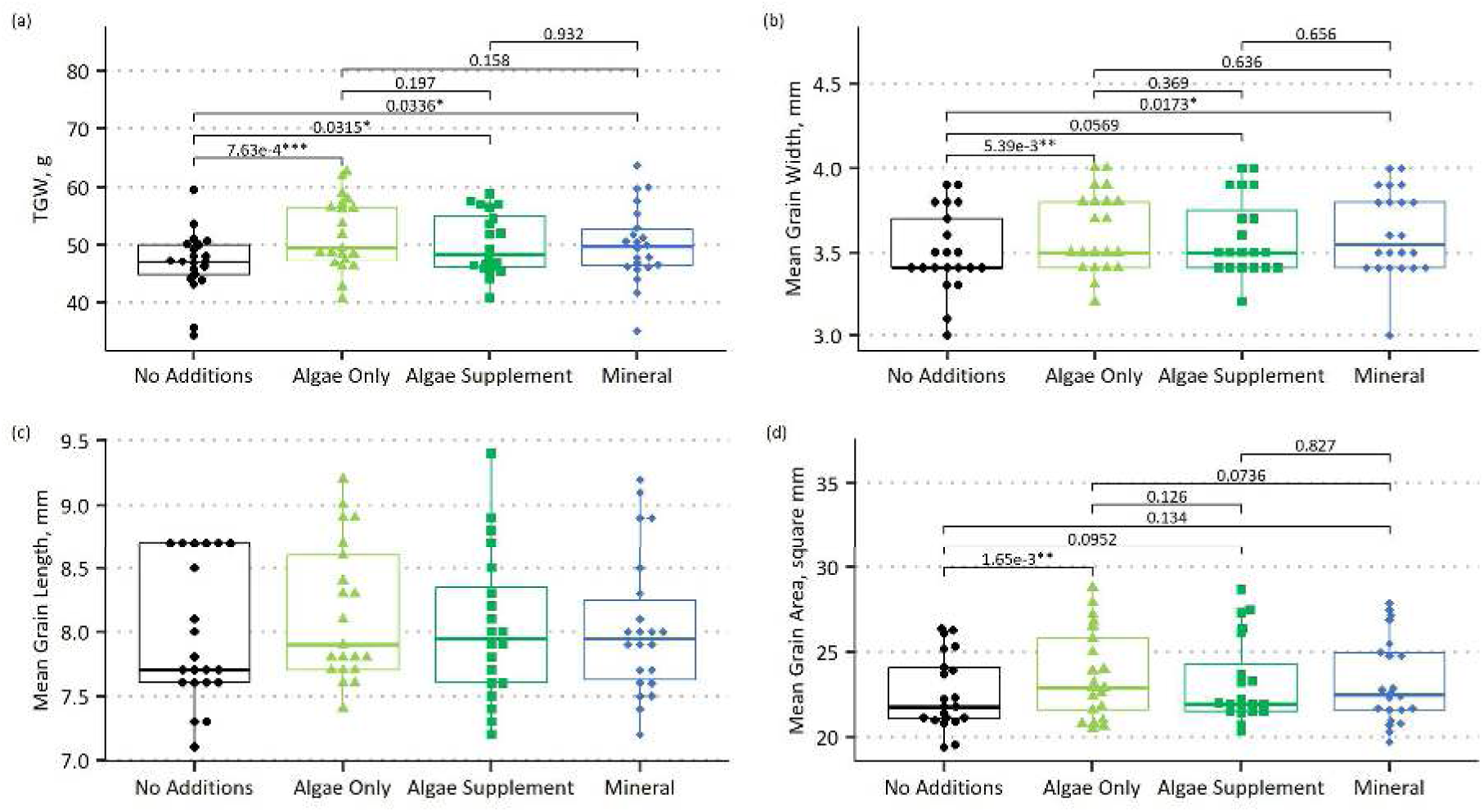
Grain traits of cv. Laureate when grown with four different nutrient sources across three glasshouse experiments. **(a)** Thousand grain weight (TGW). **(b)** Mean grain width per plant. **(c)** Mean grain length per plant. **(d)** Mean grain area per plant. For all boxplots, the thick horizontal line is the median, thin horizontal lines are the 1st and 3rd quartile. Whiskers extend to the range, excluding outliers (>1.5x the interquartile range below quartile 1 or above quartile 3). Overall significance determined by mixed model ANOVA, p-value reported in the main text. Pairwise significance determined by mixed model contrasts. N = 21 for no additions, 21 for algae only, 20 for algae supplement, 22 for mineral. Each rep is one data point

### 3.2 Spring 2024 Field Trial

#### 3.2.1 Phenology

At 26DAS, all field trial plots had 99% or 100% emergence of seedlings and were therefore considered to be fully germinated. Treatment was a significant factor for days to flag leaf unfurling (Kruskal-Wallis, p = 0.010), with plants grown with the microalgae treatment reaching flag leaf unfurling significantly sooner (median 59DAS, range 59-61) than those plants grown with no additions (median 61DAS, range 59-63), with plants grown on mineral fertiliser (median 59DAS, range 59-61) being a non-significant intermediate (**Table 2**, Online Resource 1, **Fig. S7 (a)**). For days to heading the effect of treatment is significant (Kruskal-Wallis, p = 7.82e-5), there is no significant difference between plants grown with microalgae (median 64DAS, range = 63-64) or mineral fertiliser (median 63DAS, range = 63 only), with plants grown with either treatment reaching this stage significantly sooner than plants grown with no additions (median 65DAS, range = 64-65, **Table 2**, Online Resource 1, **Fig. S7 (b)**).

**Table 2.**
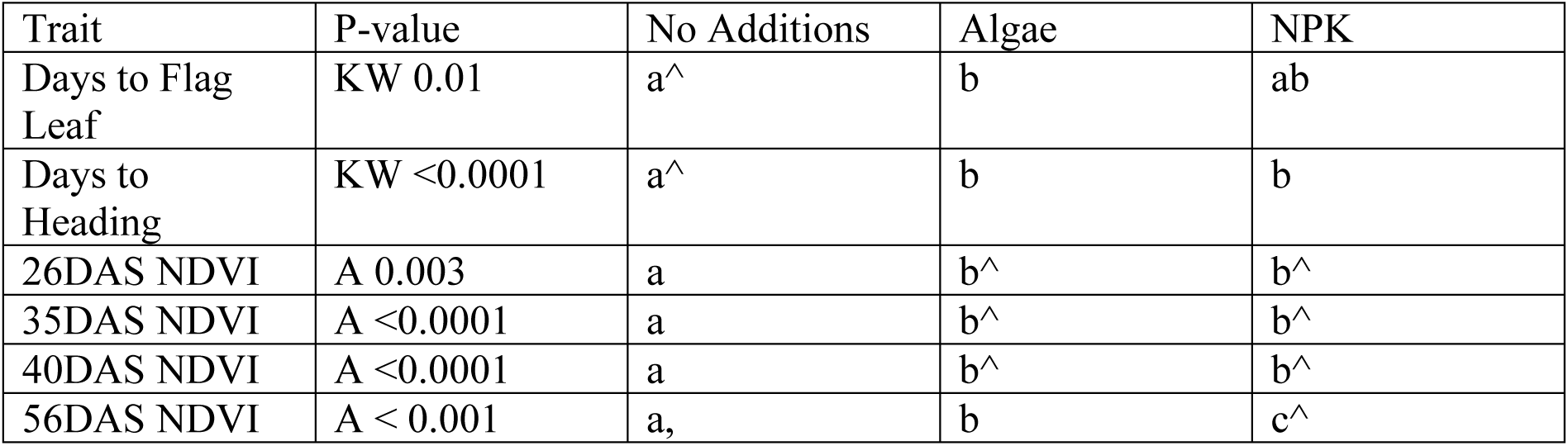
Significance of comparisons for the field trial in Online Resource 1. Graphs for each trait listed can be found in Online Resource 1, with figure numbers listed in the sections 3.2.1 and 3.2.2. For methods of obtaining p-values, KW = Kruskal-Wallis followed by Dunn Test, A = ANOVA followed by Tukey’s HSD. Different letters represent significance groups, with multiple letters representing non-significant intermediates (*i.e.* ab is not significantly different from a or b, which are significantly different from one another). The “^” sign represents the highest value in the comparison (*e.g.* No Additions has a higher Days to Flag Leaf than Algae), the “,” sign represents the lowest value in the comparison, only used when there are more than two significance groups

| Trait | P-value | No Additions | Algae | NPK |
| --- | --- | --- | --- | --- |
| Days to Flag Leaf | KW 0.01 | a^ | b | ab |
| Days to Heading | KW <0.0001 | a^ | b | b |
| 26DAS NDVI | A 0.003 | a | b^ | b^ |
| 35DAS NDVI | A <0.0001 | a | b^ | b^ |
| 40DAS NDVI | A <0.0001 | a | b^ | b^ |
| 56DAS NDVI | A < 0.001 | a, | b | c^ |

#### 3.2.2 Reflectance Indices Across the Growing Season

At 26DAS, treatment had a significant effect on NDVI (ANOVA, p = 0.0026). NDVI was not significantly different between plots supplied with microalgae or mineral fertiliser, and both treatments had significantly higher NDVIs than plants grown in plots with no additions (**Table 2**, Online Resource 1, **Fig. S8 (a)**). This suggests that plants grown with microalgae or mineral fertiliser had higher levels of leaf greenness and a better ability to absorb photosynthetically active red light. At 35DAS (ANOVA for treatment, p = 2.20e-8), NDVI had increased for the microalgae and mineral treated plots, while staying similar to the 26DAS value for the no additions plots, keeping the same relationship between treatments but increasing the level of significance (**Table 2**, Online Resource 1, **Fig. S8 (b)**). At 40DAS (ANOVA for treatment, p = 2.90e-8), NDVI starts to decrease across all treatments, which may be due to plants beginning to remobilise nutrients from the lower leaves. Nevertheless, the pattern of significant differences between treatments remains the same (**Table 2**, Online Resource 1, **Fig. S8 (c)**). At 56DAS (Mixed Model ANOVA for treatment, p < 0.001), all treatments have significantly different NDVIs from one another, with the no additions plots being the lowest, mineral being the highest and microalgae being a significantly different intermediate (**Table 2**, Online Resource 1, **Fig. S8 (d)**).

#### 3.2.3 Yield, Morphometrics and Grain Traits at Harvest

At harvest, grain moisture was between 17% and 18.9%. Grain yield, corrected to 0% moisture, was not significantly different between plots grown with microalgae or mineral fertiliser, and plots grown with either of these treatments yielded significantly more grain than plots grown with no additions (ANOVA, p < 0.0001, **Fig. 3 (a)**). Plants grown with microalgae had a statistically significantly higher number of grains per ear (median 21, range = 18-24) than those grown with mineral fertiliser (median 20, range = 17-23), with the plants grown with no additions (median 20, range = 16-25) being a non-significant intermediate (ANOVA, p = 0.023, **Fig. 3 (b)**). The plants from plots grown with microalgae also have significantly longer ears than both those grown with no additions or mineral fertiliser, which are not significantly different to each other (Mixed model ANOVA, p = 0.008, **Fig. 3 (c)**).

**Fig. 3.**
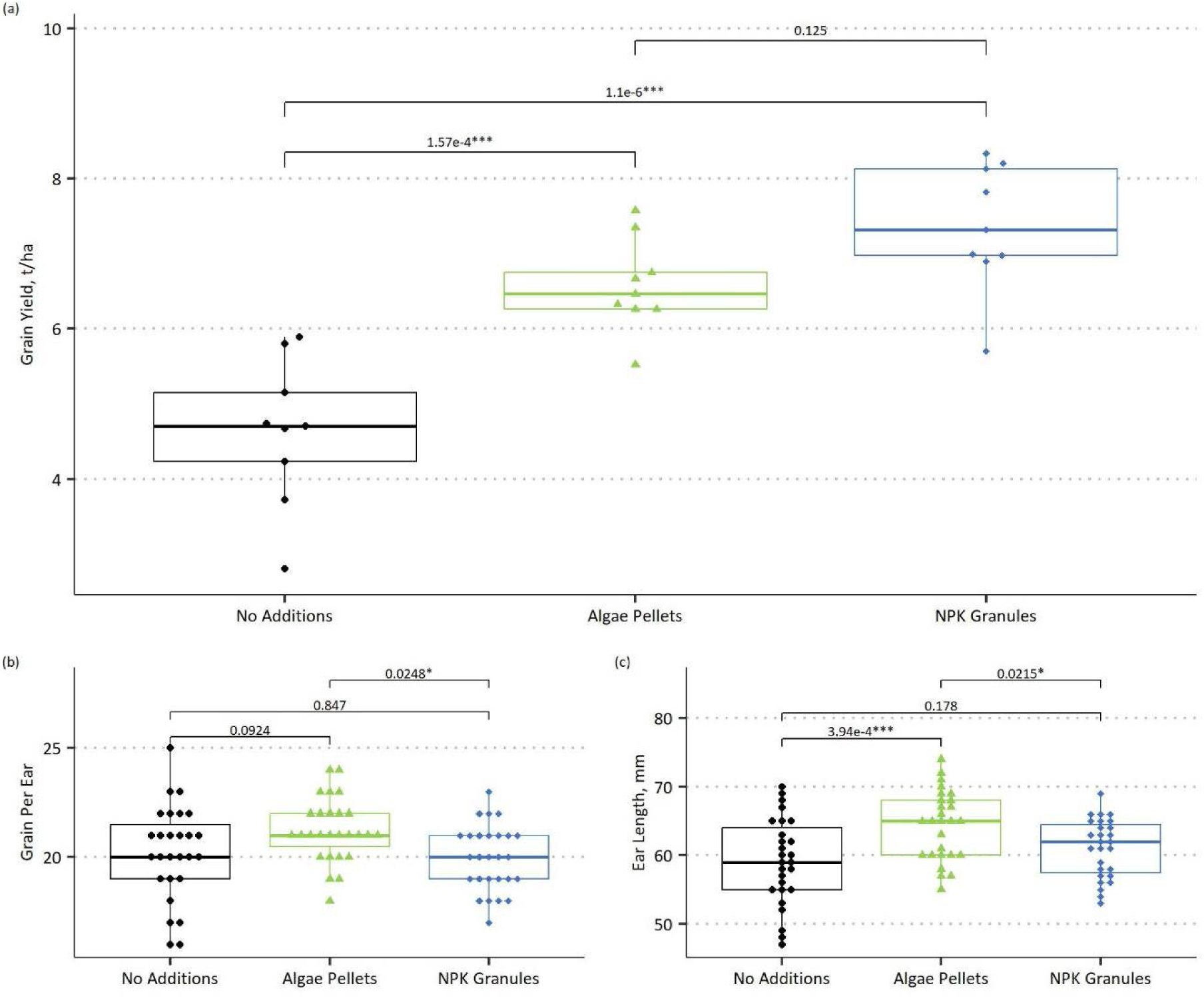
Harvest and grab sample traits of cv. Laureate when grown in a field trial under three different fertiliser regimes. **(a)** Dry grain yield per plot. **(b)** Grain per ear for three random ears in each grab sample. **(c)** Ear length for three random ears in each grab sample. For all boxplots, the thick horizontal line is the median, thin horizontal lines are the 1st and 3rd quartile. Whiskers extend to the range, excluding outliers (>1.5x the interquartile range below quartile 1 or above quartile 3). For **(a)** and **(b),** overall significance determined by ANOVA, p-value reported in main text, and pairwise significance determined by Tukey’s HSD. For **(c)**, the block effect was significant, so a mixed model was used. Overall significance determined by mixed model ANOVA, p-value reported in main text, and pairwise significance determined by mixed model contrasts. For **(a)**, N = 9 for all treatments, each rep is one point. For **(b)** and **(c)**, N = 27 for all treatments, each ear is one data point

Plots grown with microalgae or mineral fertiliser do not significantly differ in their TGW, and both have significantly higher TGWs than plots grown with no additions (ANOVA, p = 0.00036, **Fig. 4 (a)**). There is no significant difference in grain width between any treatment (**Fig. 4 (b)**). Plots grown with mineral fertiliser have significantly longer grain than those grown with no additions, with the microalgae treatment being a non-significant intermediate (Kruskal-Wallis, p = 0.0013, **Fig. 4 (c)**). This same pattern is seen for overall grain area, mineral plots having grains with significantly more area than no additions plots, with microalgae fertilised plots being a non-significant intermediate. (ANOVA, p = 0.007, **Fig. 4 (d)**).

**Fig. 4.**
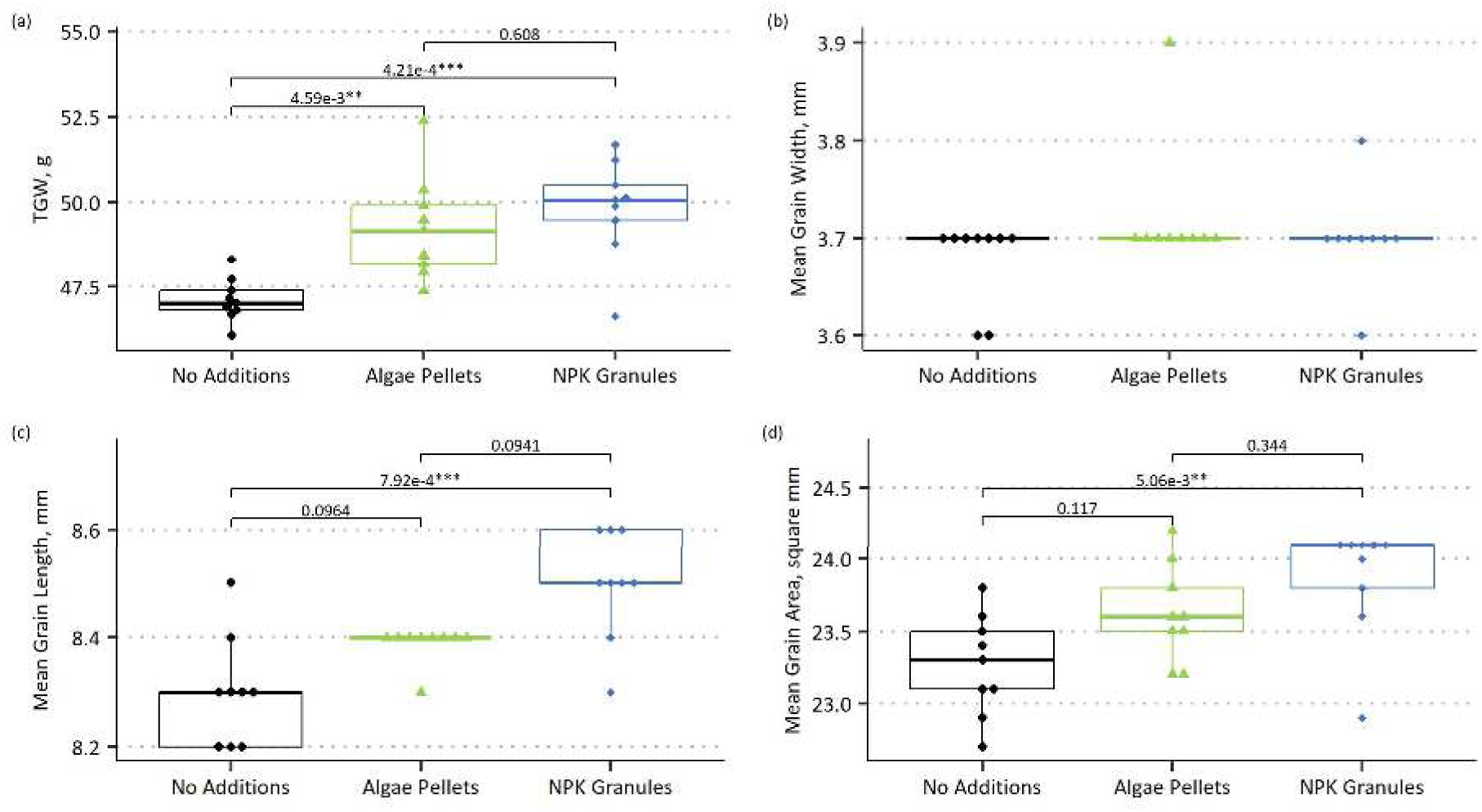
Grain mass and dimensions of cv. Laureate when grown in a field trial under three different fertiliser regimes. **(a)** Thousand grain weight on a dry basis per plot. **(b)** Mean grain width per plot. **(c)** Mean grain length per plot. **(d)** Mean grain area per plot. For all boxplots, the thick horizontal line is the median, thin horizontal lines are the 1st and 3rd quartile. Whiskers extend to the range, excluding outliers (>1.5x the interquartile range below quartile 1 or above quartile 3). For **(a)** and **(d),** overall significance determined by ANOVA, p-value reported in main text, and pairwise significance determined by Tukey’s HSD. For **(b)**, no p-value was determined. For **(c)**, overall significance determined by Kruskal-Wallis, p-value reported in main text. Pairwise significance determined by Dunn test with Benjamini-Hochberg correction for multiple testing. N = 9 for all treatments. Each rep is one data point

#### 3.2.4 Grain Quality and Malting

Grain nitrogen on a dry basis was not significantly different between plants grown with microalgae or no additions, while plants grown with mineral fertiliser had significantly higher grain nitrogen than both other treatments (Mixed model Anova, p = 0.004, **Fig. 5 (a)**). When malted, all three treatments have significantly different total malt nitrogen, with the malt from plots grown with no additions having the lowest, malt from mineral supplied plots having the highest and malt from plots grown with microalgae being a significant intermediate (Mixed model ANOVA, p < 0.001, **Fig. 5 (b)**). For soluble nitrogen, there is no significant difference between malt from barley grown with microalgae or mineral fertiliser, and grain from plants grown with either of these fertilisers give malt with higher soluble nitrogen than malt grown with no additions (Mixed Model ANOVA, p < 0.001, **Fig. 5 (c)**). There are no significant differences in soluble nitrogen as a percentage of total nitrogen (ANOVA, p = 0.24, **Fig. 5 (d)**).

**Fig. 5.**
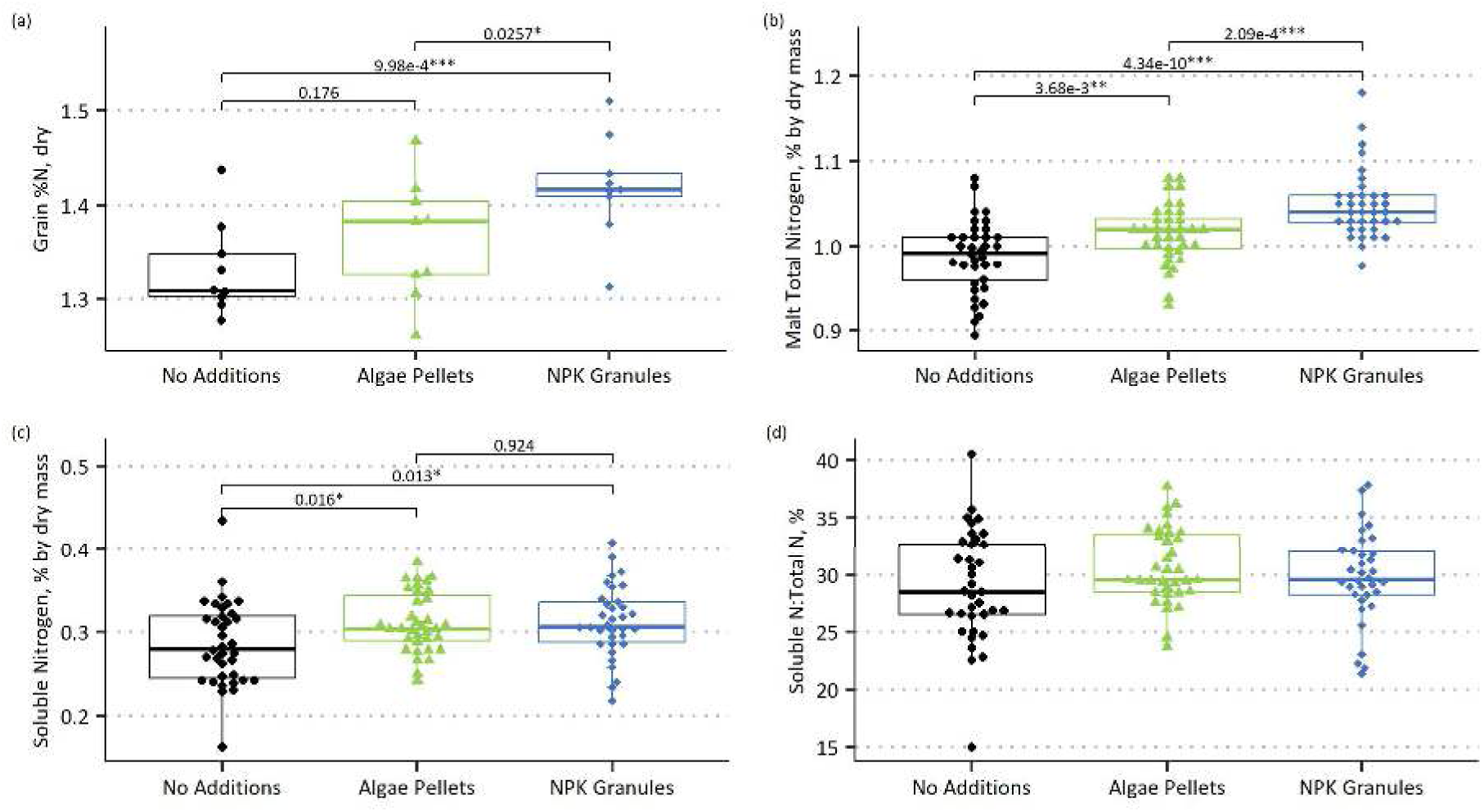
Grain and malt nitrogen of cv. Laureate when grown in a field trial under three different fertiliser regimes. **(a)** Grain %Nitrogen by dry mass. **(b)** Malt total %Nitrogen by dry mass. **(c)** Malt soluble %Nitrogen by dry mass. **(d)** Malt soluble nitrogen as a percent of malt total nitrogen. For all boxplots, the thick horizontal line is the median, thin horizontal lines are the 1st and 3rd quartile. Whiskers extend to the range, excluding outliers (>1.5x the interquartile range below quartile 1 or above quartile 3). For **(a)**, **(b)** and **(c),** overall significance determined by mixed model ANOVA, p-value reported in main text, and pairwise significance determined by mixed model contrasts. For **(d)**, significance was determined by ANOVA, p-value reported in main text. For **(a)**, N = 9 for all treatments, each point is one plot. For **(b)**, N = 36. For **(c)** and **(d)**, N = 36 for no additions, 35 for algae and 34 for mineral. Each replicate is one data point Malt moisture content ranged between 4.86% to 8.52%. Fine hot water extract strength (HWE), not moisture corrected, did not significantly differ between malt from the microalgae or mineral fertiliser treatments, and both had higher HWEs than malt from barley grown with no additions (Mixed model ANOVA, p < 0.001, **Fig. 6 (a)**). This pattern is maintained when HWE is corrected for moisture, now with increased significance due to range shrinkage (ANOVA, p < 0.0001, **Fig. 6 (b)**). There are no significant differences in predicted spirit yield (PSY) before it has been moisture corrected, as this value is highly affected by the variable malt moisture (Mixed model ANOVA, p = 0.061, **Fig. 6 (c)**). After correcting for moisture content, there is no significant difference in PSY between malt grown with microalgae or mineral fertiliser, and both malts have significantly greater PSY than malt grown with no additions (ANOVA, p = 0.002, **Fig. 6 (d)**).

**Fig. 6.**
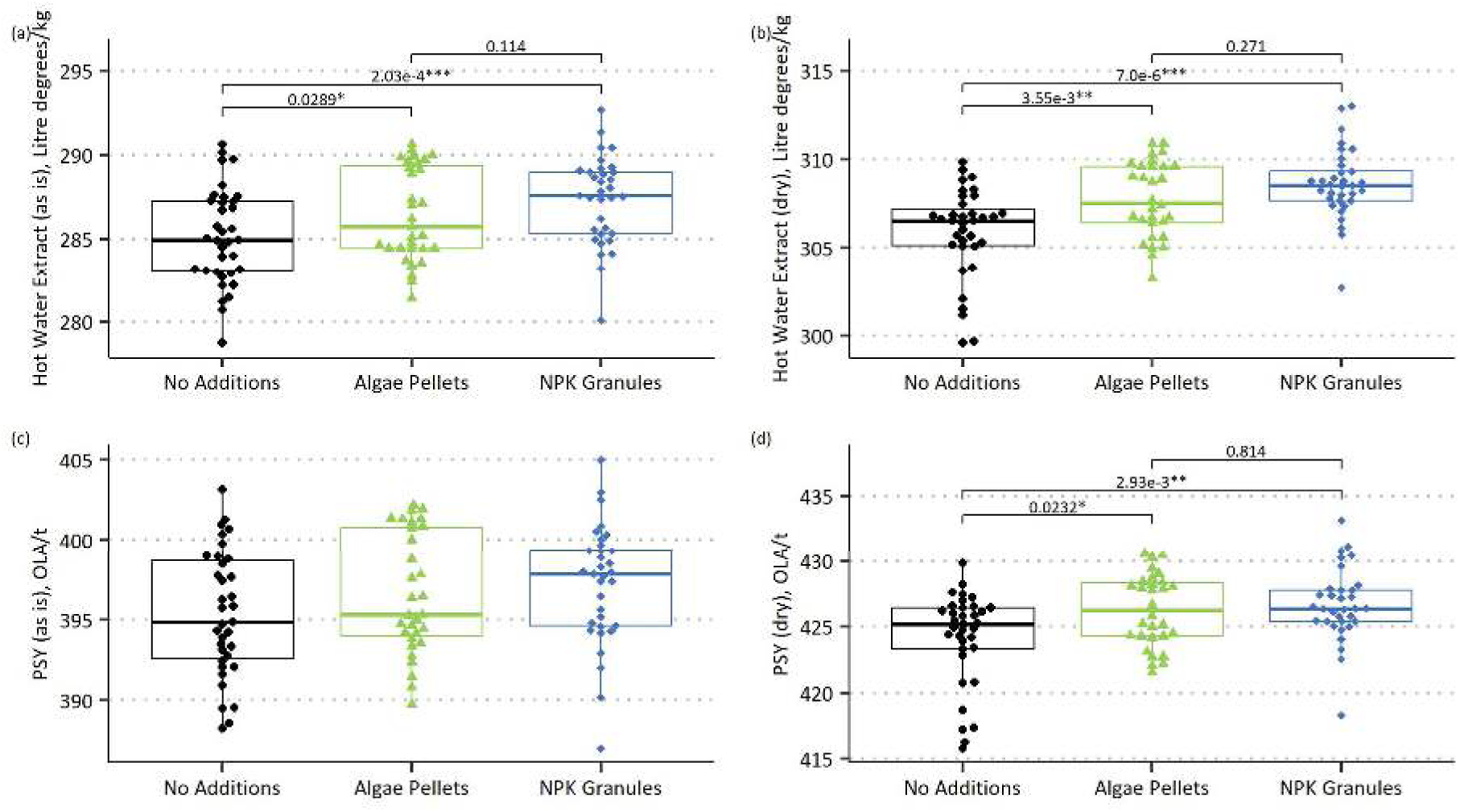
Fine hot water extract strength (HWE) and predicted spirit yield (PSY) of cv. Laureate when grown in a field trial under three different fertiliser regimes. **(a)** HWE not corrected for grain moisture. **(b)** HWE corrected to 0% grain moisture. **(c)** PSY not corrected for grain moisture. **(d)** PSY corrected to 0% grain moisture. For all boxplots, the thick horizontal line is the median, thin horizontal lines are the 1st and 3rd quartile. Whiskers extend to the range, excluding outliers (>1.5x the interquartile range below quartile 1 or above quartile 3). For **(a)** and **(c),** overall significance determined by mixed model ANOVA, p-values reported in main text, and pairwise significance determined by mixed model contrasts. For **(b)** and **(d)**, significance was determined by ANOVA, p-values reported in main text, followed by Tukey’s HSD. N = 35 for no additions, 31 (**(a)** and **(c)**)/30 (**(b)** and **(d)**) for algae and 33 for mineral. Each data point is one replicate

## 4 Discussion

In this study we have demonstrated that there is no metric related to NDVI or yield components in which barley cv. Laureate fertilised with *Chlorella vulgaris* instead of mineral fertiliser is significantly worse than barley grown with mineral fertiliser in both glasshouse and field trials, except NDVI at 56DAS for field plots. On this occasion, mean NDVI for microalgae treated plots is significantly lower than that of mineral treated plots. However, this does not seem to impact the malt quality of the grain, as there is no significant difference in fine hot water extract strength or predicted spirit yield from malt of barley grown with microalgae or mineral fertiliser.

Lima e Silva *et al*. (2024) reported an increased speed of growth of barley (experimental setting and cultivar not stated) between 40-100 days after sowing using aerial length, with increasing concentrations of *Arthrospira* sp., a cyanobacterium which is also commonly used as a protein rich health food like *C. vulgaris*. This roughly agrees with the increase in node development we saw in our first experiment between 42-46DAS, but Lima e Silva *et al*. (2024) report a much longer period of time where the plants grown with cyanobacteria show an increased growth rate, whereas we saw only a few days where the microalgae treated plants were ahead of plants grown with the other treatments, and only in one replicate of the experiment, with the other two glasshouse experiments showing no difference in phenology at any time. The minor differences in speed of development between treatments, for the first replicate of the glasshouse experiment only, and between experiments do not have an impact on the plant’s ability to use one type of fertiliser or the other, as evidenced by the lack of significant differences between the microalgae and mineral fertiliser in morphometrics or yield components.

Concerns about microalgae not releasing nutrients as quickly as mineral fertiliser and therefore starving the plant in early development (Coppens *et al*. 2015) are not supported by our results. During early tillering (GS 21-23) there are no significant differences in leaf dimensions, aboveground biomass or NDVI between plants grown with any of the treatments. This agrees with Siebers *et al*. (2019) and Mau *et al*. (2021), who both showed that P from algal sources was readily accessible and sufficient to sustain wheat growth for up to six weeks after sowing.

In three glasshouse trials, each performed at a different time of year, in soils of similar texture but different residual nutrient content, with different amounts of microalgae and mineral fertiliser added (up to the common N rate of 118kg(N)/ha), barley cv. Laureate accumulated as much biomass and developed a number of tillers that was not significantly different between plants grown with microalgae or mineral fertiliser. These results agree with the work of Schreiber *et al*. (2018) who observed that, in glasshouse trials using the wheat cultivar Scirocco grown in the soil-like substrate “Null Erde”, plant size (from projected leaf area) and shoot dry mass were not significantly different when growing with *C. vulgaris* (the same microalgal species we used here) or mineral fertiliser. Additionally, as we observed, Schreiber *et al*. (2018) reported that both treatments produced wheat plants with projected leaf areas and shoot dry masses that were significantly higher than plants grown with no additions. Our work extends beyond Schreiber *et al*. (2018), who terminated their experiment before their wheat plants reached maturity. Renuka *et al*. (2016) grew wheat plants, var. HD2967, to maturity in growth chambers, using consortia of algae made up of multiple taxa, one of which included species of *Chlorella*, to partially replace mineral fertiliser in a compost mix. Like the data presented in this study, Renuka *et al*. (2016) found that dry mass at maturity was not significantly different to, or was higher than, the full NPK control when using their algae treatments. We observed in glasshouse and field trials that the number of filled grain per plant and TGW are not significantly different between plants grown with microalgae or mineral fertiliser (and that they exceed the no additions negative control), while Renuka *et al*. (2016) found that TGW significantly increased for plants grown with consortia of algae over the full NPK control, and far exceeded the TGW of the 75%N negative control. This result is beyond what we observed, although both our findings agree that there is no reduction in TGW when microalgal fertiliser is used instead of mineral fertiliser. In contrast, the work of Mückschel *et al*. (2023) disagrees with the findings of Renuka *et al*. (2016). They observed that when using consortia of algae as fertiliser, wheat cv. KWS Starlight growing in a glasshouse in soil still gives significantly higher shoot dry masses and grain yield per plant than unfertilised controls but had significantly lower values than plants grown with mineral fertiliser, which they attribute to lower availability of the forms of nitrogen in microalgae compared to mineral forms. While this wasn’t seen in the results of this paper, the work of Schreiber *et al*. (2018) or Renuka *et al*. (2016), it is possible that the availability of nutrients from algal fertiliser may be influenced by the soil type or the microbiome within, showing that more extensive testing in a wider range of conditions is necessary.

The translatability between wheat and barley is positive, although not entirely unexpected, as far more distantly related crops have shown similar responses to the potential of microalgae as a fertiliser, such as tomato (Garcia-Gonzalez *et al*. 2016) or lettuce (Faheed *et al*. 2008).

Additionally, this also shows that a range of formulation methods can be employed to give similar results, with some exceptions. The current study applied algae and mineral fertiliser to provide equivalent nitrogen, whereas Schreiber *et al*. (2018) added algae and fertiliser to provide equivalent phosphorus. Renuka *et al*. (2016) added algae based on green mass (µg chlorophyll per gram of carrier), which resulted in a not statistically different amount of nitrogen being present in the positive control and algae supplemented treatments. In contrast, Mückschel *et al*. (2023) added N, P and K salts to match the amounts of these elements found in their algae consortia. This implies that, *if no nutrient is lacking in amount or availability* in a certain formulation, microalgae can be used to wholly or partially substitute for either N or P, and likely any other nutrient, provided that the nutrients in that microalgae can be made available to the plant.

The importance of understanding the nutrient content of microalgal fertiliser and applying it at suitable rates, comparable to what one would use for mineral fertiliser, is demonstrated by the work of Alobwede *et al*. (2019). They found that when adding *Chlorella* sp. based on mass of algae per area, not mass of individual nutrient per area, to provide 24.28kg(N)/ha to field plots planted with wheat cv. Tybalt, there was no significant difference in shoot dry mass or ear number relative to the no algae control. As hypothesised in their discussion, they likely used too low a mass of algae to find a significantly different response, although without a mineral fertiliser positive control, it may be the case that their soil was replete with nitrogen such that adding additional nitrogen would give no increase in plant performance. In contrast, our study added 61.6kg(N)/ha of microalgae or mineral fertiliser to our field trial in addition to the baseline soil nitrogen and did find a significant increase in yield and yield components of barley over the no additions negative control, and no significant difference in yield between the microalgae and mineral fertiliser treatments.

The increase in yield in the microalgae and mineral fertiliser treatments over the negative control, in this study, appears to be driven by a combination of increased TGW, as seen for both the glasshouse and field trials, and increased tiller number leading to an increase in total filled grain per plant, seen in the glasshouse experiments. Grain per ear does not significantly differ between treatments in the glasshouse experiment, and the medians only differ by one grain in the field trial and therefore does not contribute to an increase in overall grain yield per plant or per plot. The agreement of our glasshouse and field trials regarding yield and most yield components is positive, although there are some differences in which treatments are significantly different in phenology, NDVI and grain dimensions between the glasshouse and field trials. While these differences did not affect overall yield and our conclusions are the same, more field trials in a wider range of soils and local climates with more frequent monitoring of NDVI would be valuable to assess if microalgae can consistently support the ability of the plant to absorb photosynthetically active red light later in development.

As barley is a key component of the high value markets of brewing and distilling, the examination of malt quality was a necessary and novel step for this study. For grain nitrogen, our values for all treatments are well below a typical industry maximum of 1.60%, and below the target range of 1.35-1.55% (Chivas Brothers, personal correspondence). Malt nitrogen is similar, with every value being below the specification range. The significant difference in grain and malt %N between microalgae and mineral fertiliser differs from the previous pattern of yield and yield components, possibly indicating that the plant will allocate nitrogen from each source differently. While, in this study, values for grain and malt %N were below specification, the industry typically prefers the lower end of the target range for malt whisky distilling. If microalgal fertiliser consistently results in lower grain or malt %N than mineral fertiliser, while maintaining yield and other malt quality traits, this may be a boon for the users of low nitrogen malt. There is no difference in soluble nitrogen ratio (SNR) between any of the treatments, and it is below specification for most samples. SNR is a proxy for malt modification (the extent of mobilisation of the sugars and proteins within the endosperm during germination), with a higher value indicating more modification. The low values found for SNR in this study imply under-modification during the malting process, which in our case is likely due to poor germination rather than insufficient germination time. Extract strength as-is and dry extract strength for microalgae and mineral fertiliser were not significantly different from each other, and both had significantly greater extract strength than malt grown with no additions. However, dry extract strength is again below the specification minimum for all treatments. Predicted spirit yield (PSY) as-is is not significantly different between treatments, which can be explained by the wide distribution of malt moistures. Accounting for moisture content, dry PSY again agrees with the earlier measurements, where microalgae and mineral fertiliser are not significantly different in PSY, and have significantly higher PSYs then malt grown with no additions. In this case, the dry PSY for all treatments meets the specification. Despite extract strength, which is proportional to PSY, being below specification, the low %N, which is inversely proportional to PSY, increases the PSY to an acceptable value. However, the PSY calculation may only be valid for %N in the range specified, and too low a %N may impact the ability of starch in the grain to be mobilised during malting and mashing due to lack of enzymes, or lack of free amino nitrogen, which yeast relies on for growth during fermentation (Chivas Brothers, personal correspondence).

The low values for grain and malt %N for all treatments may be due to a combination of two factors. Firstly, the entirety of the fertiliser was applied at the start of the growing period. On some farms, the timing of application of fertiliser has been reported to change the grain %N, with earlier applications decreasing grain %N, and later applications (*e.g*. as a top dressing) increasing grain %N (Kendall *et al*. 2021). Secondly, the trial area was at the top of a sloping field. Based on weather station data from the James Hutton Institute, Dundee, a short way from Balruddery farm where this field trial was conducted, May of 2024 had ∼20mm more rainfall than the 30 years average (70.8mm in 2024 compared to 51.2mm 1991-2020). The increased rainfall may have led to more of the residual soil N and applied N being washed away during the spring, decreasing the nitrogen available to the plant and thus decreasing the grain %N. To provide some N later in the growing period and avoid the potential losses from late-spring rain, part of the total fertiliser may be added later, for example at growth stage 31-33 (early stem elongation), a common stage for a second application. However, if microalgae is applied as pellets on the soil surface at this stage, it may face difficulty with incorporation into soil and therefore reduce nutrient availability to the plant. This could be mitigated by having an initial ploughing-in of algae at sowing, making up most of the total nutrient load, followed by an application of the more soluble mineral fertiliser at a later growth stage, although this has the obvious disadvantage of still requiring some mineral fertiliser. Field trials in different locations over multiple years with different fertiliser splits, supplemented by regular soil testing across the growing season to determine the rate of mineralisation (and therefore availability) of microalgae provided N, are needed to properly assess the viability of use of microalgae as a fertiliser for malting barley, but this study is a promising start given that microalgae and mineral fertiliser did not differ in terms of extract and PSY.

Unfortunately, microalgal fertilisers currently have barriers to widespread acceptance. Firstly, they are not economically viable enough to outcompete mineral fertiliser, if the microalgae is cultivated solely for fertiliser. However, charging a premium for microalgae grown produce, as is already done for produce grown with other organic fertilisers, or using microalgae as wastewater treatment, reducing the cost through turning a waste product into a saleable co-product, could be solutions to this problem (Coppens *et al*. 2015; Ronga *et al*. 2019).

Recently, there was a world fertiliser shortage due to geopolitical events increasing the cost of energy and increasing difficulties in shipping (Polanesk *et al*. 2022). Short-term mineral fertiliser shortages and long-term sustainability goals could incentivise both farmers and the biotechnology sector into adopting more sustainable solutions, such as microalgae derived products including microalgal fertiliser. Secondly and applying only to wastewater grown microalgae, depending on the type of wastewater used there may be accumulation of heavy metals, pharmaceuticals, or other toxic compounds. Any algal product grown on wastewater at risk of accumulating these compounds should be tested thoroughly to ensure they are safe and suitable for use in agriculture, with concentrations of harmful compounds below the limits imposed by local authorities, as is already done for other organic fertilisers in Scotland (Farm Advisory Service 2021).

In conclusion, this study demonstrates that barley cv. Laureate can use microalgal fertilisers as effectively as mineral fertilisers, in both glasshouse and field settings, within the context of the phenological, morphometric and yield components analysed in this study. There is no penalty to use of microalgal fertiliser in any measured metric related to morphometrics or yield, and significant improvements over the unfertilised, negative control were seen for microalgal and mineral fertiliser when examining yield and yield components. Malting quality was below industry specification for all treatments, but within the experiment, malt from plants grown with microalgae performed the same as malt grown with the recommended rate of mineral fertiliser, except for total malt nitrogen. Our results give support for further experiments, both in controlled and field conditions, to allow for investigation into molecular mechanisms, genotypic variation in response, economic and environmental feasibility, and additional trials using microalgae specifically grown on distillery co-products and waste to assess the practicality of nutrient recycling within the barley to malt to whisky value chain.

## Author Contributions

DJA: Conceptualization, data curation, formal analysis, investigation, methodology, visualization, writing – original draft, writing – review & editing

KH: Conceptualization, funding acquisition, supervision, writing – review & editing

DB: Resources, supervision, writing – review & editing

TM: Funding acquisition, writing – review & editing

SM: Resources, methodology, writing – review & editing

## Data Availability

The datasets generated for this study can be found at Zenodo.org under ‘Datasets supporting the paper “Barley Can Utilise Algal Fertiliser to Maintain Yield and Malt Quality Compared to Mineral Fertiliser“’ at https://doi.org/10.5281/zenodo.16810830.

## Acknowledgements

We would like to acknowledge the assistance of Richard Keith, Christopher Warden and Derek Matthews for their assistance with organising, planting, treating, and harvesting the field trial. We would also like to thank Jim Wilde and Alison Dobson as glasshouse technicians, Niki McCallum as lab manager, and Cameron McCarthy at Chivas Brothers, Glen Keith Technical Centre for overseeing David Ashworth’s malt analysis competency training.

