## Supplementary Material for "Barley Can Utilise Algal Fertiliser to Maintain Yield and Malt Quality Compared to Mineral Fertiliser"

For

Published in

**Plant and Soil**

**David James Ashworth**[1,2] (0009-0009-3687-8257), **Stefan Masson**[3], **Tom Mulholland**[3], **Davide Bulgarelli**[2] (0000-0002-2020-6642), **Kelly Houston**[1]* (0000-0003-2136-245X)

[1]The James Hutton Institute, Cell and Molecular Sciences, Invergowrie, United Kingdom

[2]University of Dundee, Division of Plant Sciences, Dundee, United Kingdom

[3]Chivas Brothers – Pernod Ricard, Dumbarton, United Kingdom

***Correspondence:**
Dr Kelly Houston


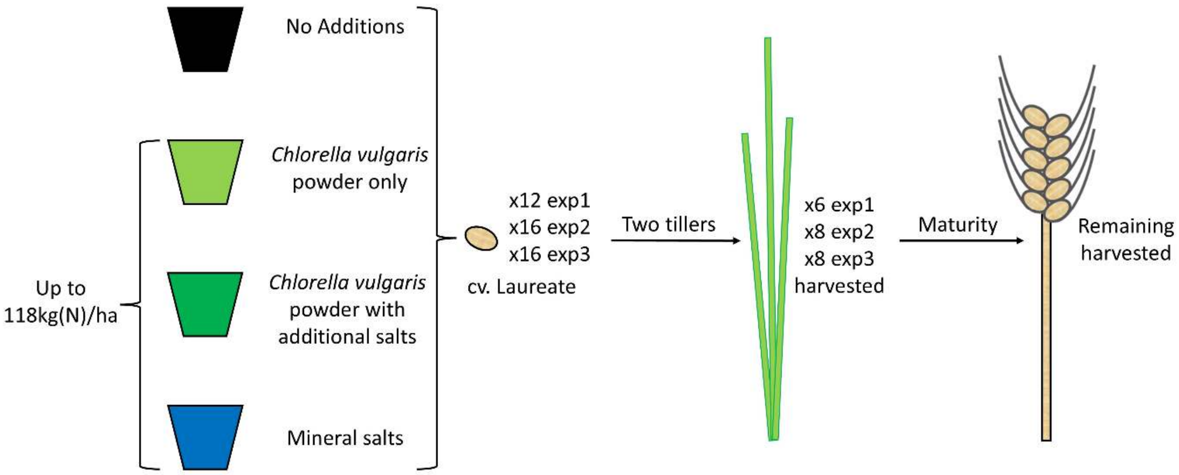

**Fig. S1 Visual summary of the treatments and harvesting points of the three replicate glasshouse experiments.** Details of each treatment, experimental design and glasshouse conditions can be found in the main text

**Table S1 Macro- and micronutrient content of the soils used for glasshouse and field experiments.** Analysis provided by Yara UK using their SA10 and BSE SOL suite of tests on a 500g sample of soil that had been passed through a 4mm sieve to remove stones

| Element | Soil | | | |
|---|---|---|---|---|
| | Bullionfield | Quarryfield 4 | Quarryfield 5 | Hutchens |
| N (mg/kg) | 54.2 | 13.6 | 30.4 | 4.8 |
| P (ppm) | 65 | 94 | 54 | 63 |
| K (ppm) | 226 | 213 | 269 | 221 |
| Mg (ppm) | 195 | 127 | 207 | 283 |
| Ca (ppm) | 1604 | 2996 | 1681 | 2155 |
| S (ppm) | 10 | 7 | 7 | 5 |
| Na (ppm) | 22 | 21 | 18 | 26 |
| B (ppm) | 1.03 | 1.49 | 0.99 | 1.16 |
| Cu (ppm) | 14.2 | 22.5 | 12.4 | 9.9 |
| Fe (ppm) | 1090 | 873 | 1079 | 665 |
| Mn (ppm) | 30 | 16 | 24 | 22 |
| Mo (ppm) | 0.05 | 0.02 | 0.02 | 0.06 |
| Zn (ppm) | 5.2 | 22.2 | 5.3 | 5.7 |

**Table S2 Composition of modified Hoagland solution.** All components except for the micronutrient stock were added as salts in the soil at the start of the experiment, in the molar ratios presented here. The micronutrient solution was added from week 4 onwards. All salts were anhydrous, except for those with waters of hydration listed

| Component | Concentration | Unit |
|---|---|---|
| $(NH_4)_2SO_4$ | 0.025 | mol/L |
| $Ca(NO_3)_2.4H_2O$ | 0.04 | mol/L |
| $KNO_3$ | 0.01 | mol/L |
| $MgSO_4$ | 0.003 | mol/L |
| FeNaEDTA | 0.0001 | mol/L |
| $KH_2PO_4$ | 0.001 | mol/L |
| 1000x Micronutrient stock | 1 | mL/L |
| 1000x Micronutrient stock composition | | |
| $MnCl_2$ | 1.188 | g/L |
| $H_3BO_3$ | 1.422 | g/L |
| $ZnCl_2$ | 0.098 | g/L |
| $CuSO_4.5H_2O$ | 0.398 | g/L |
| $Na_2MoO_4$ | 0.242 | g/L |
| $CoCl_2.6H_2O$ | 0.238 | g/L |

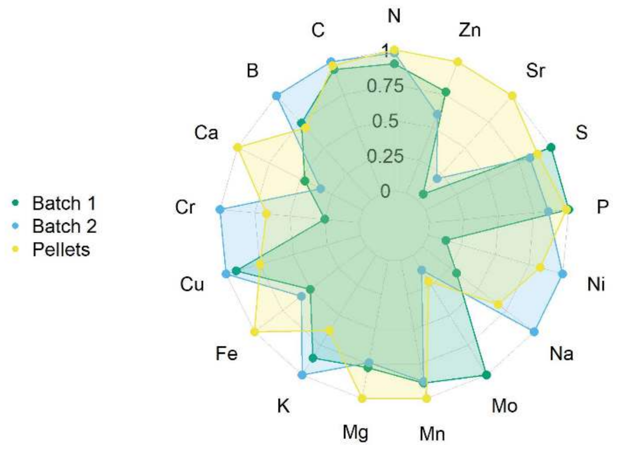

**Fig. S2 Radar plot of macronutrients, micronutrients, and heavy metals in three batches of *Chlorella vulgaris* from Raw Living Ltd.** Points are a mean of three technical replicates. The axis represents a fraction of the batch with the maximum value for that element. Exact values can be found in **Table S3**

**Table S3 Macro and micronutrient content of the algae sources used.** Analysis provided by the James Hutton Institute, Aberdeen, on a CN analyser for carbon and nitrogen, or by ICP-OES for the other elements. Values are the mean of three technical replicates. A value with a ‘less than’ sign (<) indicates that element was below the detection limit. Values presented as a range indicate at least one of the technical replicates was below the detection limit. Letters indicate significance groups, such that each different letter shows a significant difference at $p < 0.05$. Significant differences could not be determined where a technical replicate was below the detection limit

| Element | Algae Source | | |
|---|---|---|---|
| | Powder Batch 1 | Powder Batch 2 | Pellets |
| N (%dry mass) | 9.17 *a* | 9.97 *b* | 10.17 *c* |
| C (%dry mass) | 48.2 *a* | 51.1 *b* | 49.6 *c* |
| As (mg/kg) | <0.05 | <0.05 | <0.05 |
| B (mg/kg) | 8.96 *a* | 12.16 *b* | 8.41 *a* |
| Ca (mg/kg) | 896.0 *a* | 657.5 *b* | 1935.5 *c* |
| Cd (mg/kg) | <0.04 | <0.04 | <0.04 - 0.0442 |
| Co (mg/kg) | <0.05 | 0.0842 | 1.001 |
| Cr (mg/kg) | 0.532 *a* | 2.136 *b* | 1.425 *c* |
| Cu (mg/kg) | 3.20 *a* | 3.45 *b* | 2.59 *c* |
| Fe (mg/kg) | 175.1 *a* | 202.6 *a* | 348.3 *b* |
| K (mg/kg) | 6311 *a* | 7364 *b* | 4629 *c* |
| Mg (mg/kg) | 3008 *a* | 2875 *a* | 3876 *b* |
| Mn (mg/kg) | 50.1 *a* | 49.4 *a* | 56.3 *b* |
| Mo (mg/kg) | 1.050 *a* | 0.128 *b* | 0.226 *c* |
| Na (mg/kg) | 1570 *a* | 5101 *b* | 3456 *c* |
| Ni (mg/kg) | 0.163 *a* | 1.242 *b* | 1.038 *c* |
| P (mg/kg) | 11822 *a* | 10106 *b* | 11637 *a* |
| Pb (mg/kg) | <0.4 | <0.4 | <0.4 |
| Rb (mg/kg) | <5 | <5 | <5 - 8.1 |
| S (mg/kg) | 6877 *a* | 5721 *b* | 6147 *c* |
| Se (mg/kg) | 1.007 | 1.906 | <0.5 - 0.737 |
| Sr (mg/kg) | 0.617 *a* | 2.330 *b* | 11.476 *c* |
| Ti (mg/kg) | <0.02 - 2.99 | <0.02 - 1.05 | 2.01 |
| Zn (mg/kg) | 13.6 *a* | 10.6 *b* | 17.7 *c* |

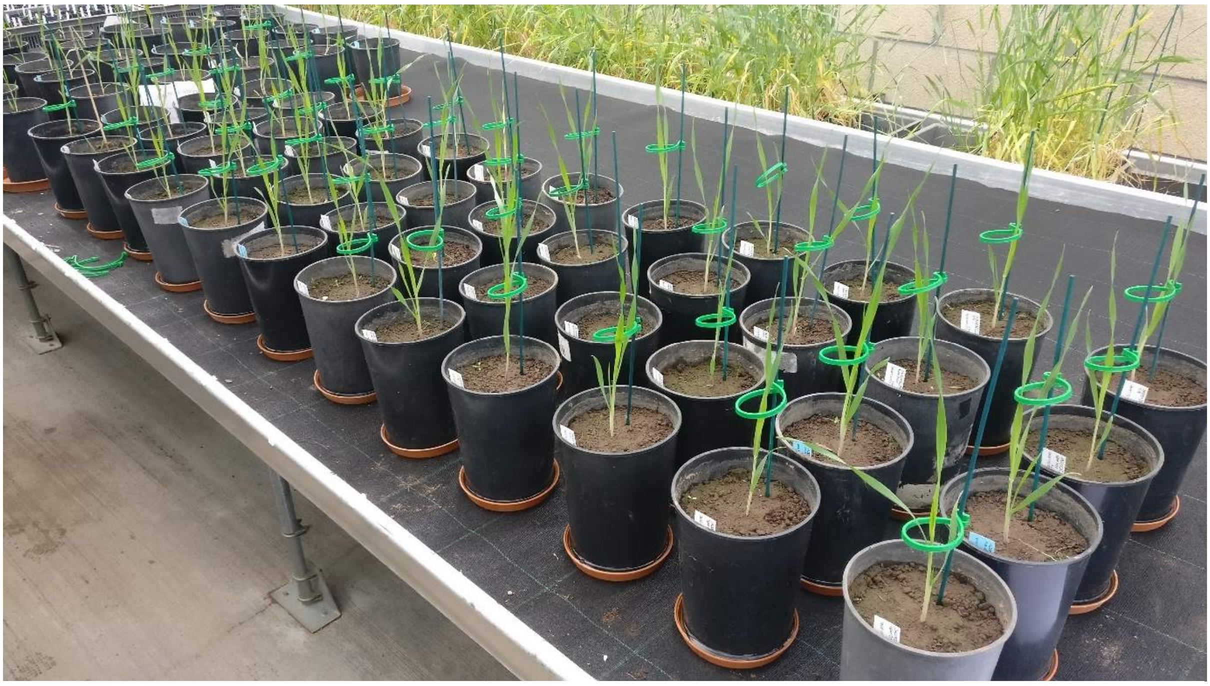

**Picture S1 Plants for the 1st Algae Response Experiment.** Pictured at the early stage of growth (GS10-19). Plants were paired by treatment, arranged in six random complete blocks and laid out on a bench in a glasshouse at the James Hutton Institute, Dundee. Picture taken by David Ashworth

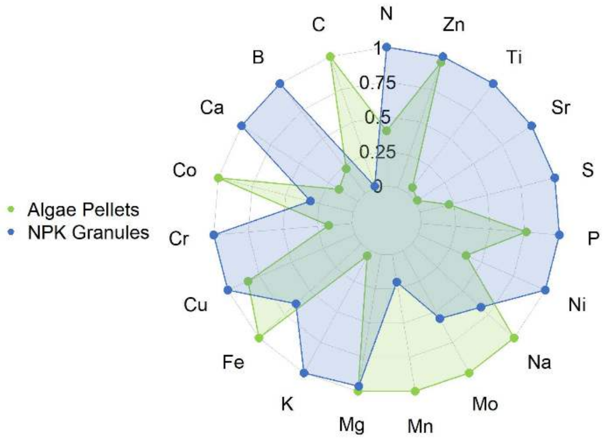

**Fig. S3 Radar plot of macronutrients, micronutrients, and heavy metals in *Chlorella vulgaris* pellets from Raw Living Ltd and 24-4-14 NPK granules.** Points are a mean of three technical replicates. The axis represents a fraction of the batch with the maximum value for that element. Exact values can be found in **Table S4**

| Element | Source | | P Value |
|---|---|---|---|
| | Algae Pellets | NPK Granules | |
| N (%dry mass) | 10.17 | 25.61 | <0.0001 |
| C (%dry mass) | 49.6 | 0.516 | <0.0001 |
| As (mg/kg) | <0.05 | 0.242 | ND |
| B (mg/kg) | 8.407 | 37.78 | <0.0001 |
| Ca (mg/kg) | 1935 | 12068 | 0.001 |
| Cd (mg/kg) | <0.04 - 0.044 | 0.268 | ND |
| Co (mg/kg) | 1.001 | 0.316 | <0.0001 |
| Cr (mg/kg) | 1.42 | 8.43 | <0.0001 |
| Cu (mg/kg) | 2.59 | 3.08 | 0.004 |
| Fe (mg/kg) | 348.3 | 222.6 | 0.002 |
| K (mg/kg) | 4629 | 111263 | <0.0001 |
| Mg (mg/kg) | 3876 | 3728 | 0.57 (ns) |
| Mn (mg/kg) | 56.3 | 11.3 | <0.0001 |
| Mo (mg/kg) | 0.226 | 0.125 | 0.002 |
| Na (mg/kg) | 3456 | 2330 | <0.0001 |
| Ni (mg/kg) | 1.04 | 2.75 | 0.00014 |
| P (mg/kg) | 11637 | 15326 | 0.002 |
| Pb (mg/kg) | <0.4 | 2.36 | ND |
| Rb (mg/kg) | <5 - 8.07 | 26.73 | ND |
| S (mg/kg) | 6147 | 28545 | <0.0001 |
| Se (mg/kg) | 0.737 | <0.5 | ND |
| Sr (mg/kg) | 11.5 | 732.5 | <0.0001 |
| Ti (mg/kg) | 2.01 | 37.17 | <0.0001 |
| Zn (mg/kg) | 17.7 | 18.5 | 0.019 |

**Table S4 Macro and micronutrient content of the algae pellets and NPK granules used in the field trial.** Analysis provided by the James Hutton Institute, Aberdeen. Values are the mean of three technical replicates. A value with a 'less than' sign (<) indicates that element was below the detection limit. Values presented as a range indicate at least one of the technical replicates was below the detection limit. The P value column shows the level of significance of the difference as reported by ANOVA. ND indicates the P value could not be determined, as at least one technical replicate was below the detection limit for that element. NS denotes no significant difference

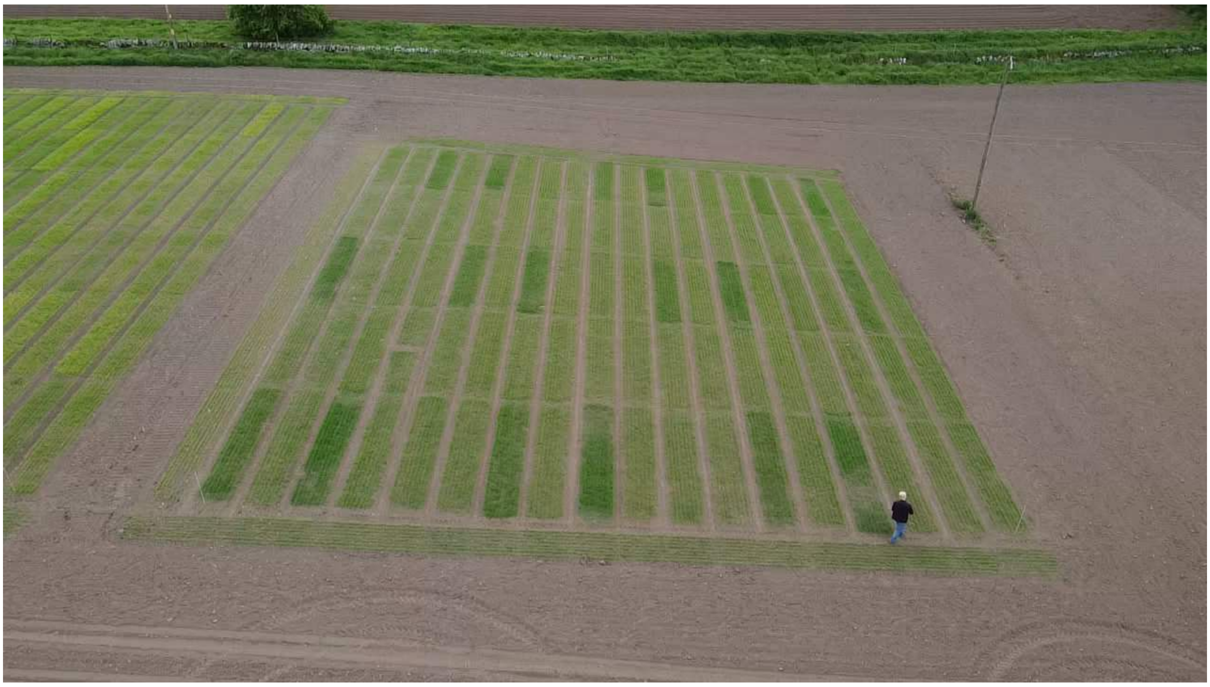

**Picture S2 Drone picture of the Spring 2024 Field Trial.** Pictured at 26 days after sowing. Experimental plots of cv. Laureate received either no additions, algae pellets or NPK granules. Treatments were distributed as 3x3 Latin Squares to account for heterogeneity of soil. Experimental plots were separated on all axes by guard plots of cv. LG Diablo. Darker plots are those treated with either algae or mineral fertiliser, lighter plots are those given no additions, or guard plots. David Ashworth (co-author) is present in the image and has given permission for his likeness to appear. Drone operated by Christopher Warden

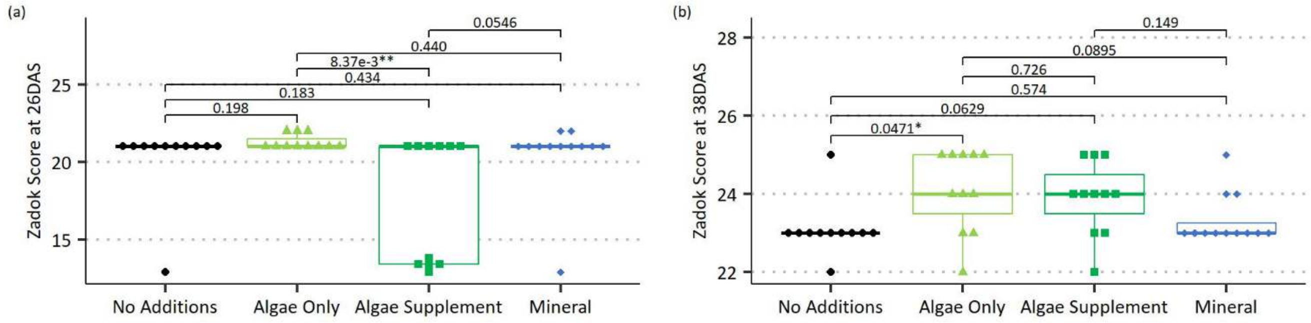

**Fig. S4 Zadoks Score of cv. Laureate in the first algae response experiment under four different nutrient treatments during tillering.** Zadok Score is a measure of plant development that categorises the major growth stages and the stages within *e.g.* GS20-29 are measures of tillering, GS30-39 are measures of shoot elongation. **(a)** Zadoks score at 26DAS. **(b)** Zadoks score at 38DAS. For both boxplots, the thick horizontal line is the median, thin horizontal lines are the 1st and 3rd quartile. Whiskers extend to the range, excluding outliers (>1.5x the interquartile range below quartile 1 or above quartile 3). Overall significance determined by the Kruskal-Wallis test, p-value reported in the main text. Pairwise significance determined by Dunn test, corrected for multiple testing using the Benjamini-Hochberg method. N = 11 for no additions, Algae Only, and Algae Supplement, N = 12 for mineral. Each rep is one data point

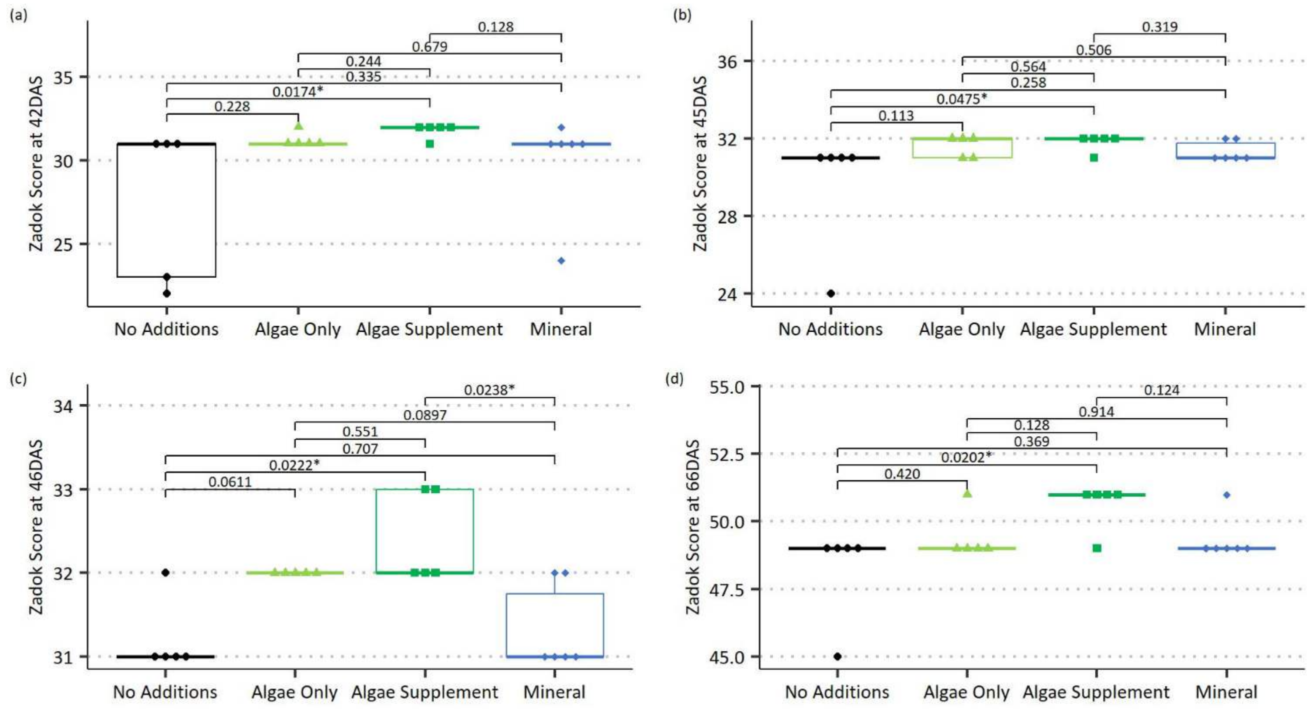

**Fig. S5 Zadoks score of cv. Laureate in the first algae response experiment under four different nutrient treatments during stem elongation and ear emergence. (a)** Zadoks score at 42DAS. **(b)** Zadoks score at 45DAS. **(c)** Zadoks score at 46DAS. **(d)** Zadoks score at 66DAS. For all boxplots, the thick horizontal line is the median, thin horizontal lines are the 1st and 3rd quartile. Whiskers extend to the range, excluding outliers (>1.5x the interquartile range below quartile 1 or above quartile 3). Overall significance determined by the Kruskal-Wallis test, p-value reported in the main text. Pairwise significance determined by Dunn test, corrected for multiple testing using the Benjamini-Hochberg method. N = 5 for no additions, Algae Only, and Algae Supplement, N = 6 for mineral. Each rep is one data point

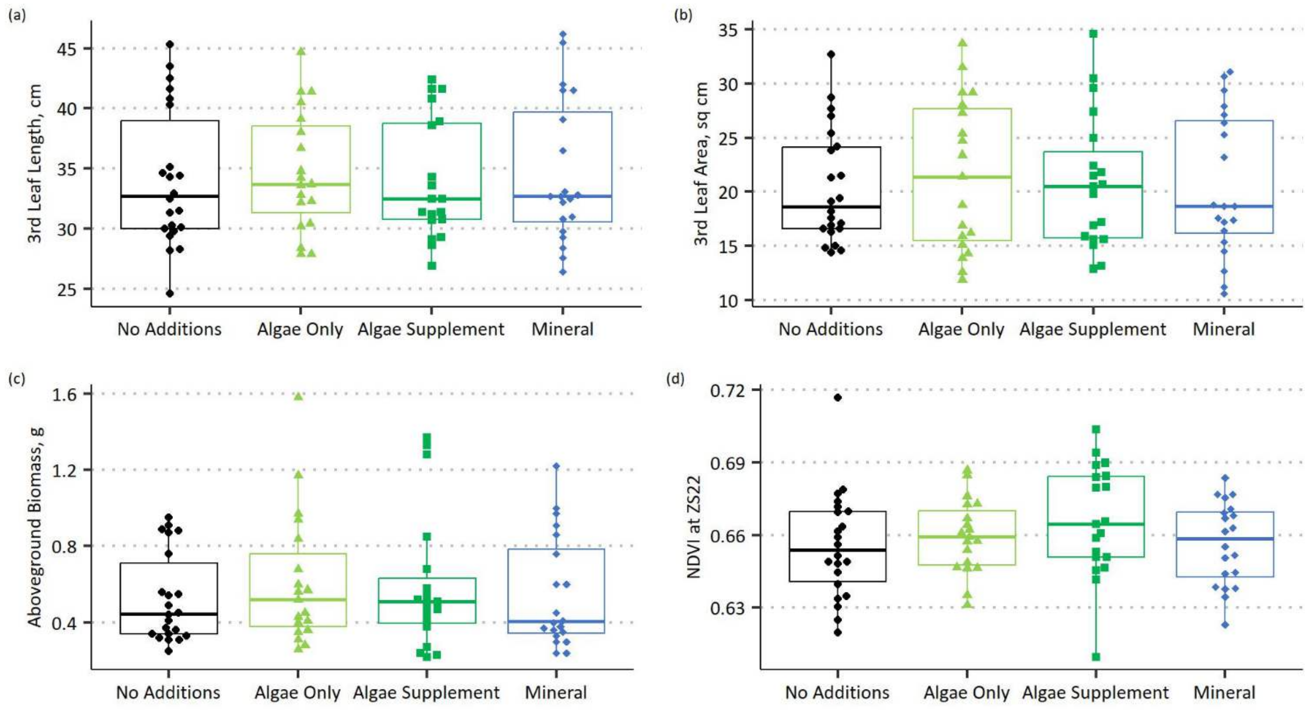

**Fig. S6 Morphometrics and reflectance of cv. Laureate at GS22 when grown with different sources of nutrients across three glasshouse experiments. (a)** Third leaf length. **(b)** Third leaf area. **(c)** Aboveground dry biomass. **(d)** NDVI of the third leaf. For all boxplots, the thick horizontal line is the median, thin horizontal lines are the 1st and 3rd quartile. Whiskers extend to the range, excluding outliers (>1.5x the interquartile range below quartile 1 or above quartile 3). Overall significance determined by mixed model ANOVA, p-value reported in the main text. No pairwise comparisons were necessary. For **(c)**, the model fitted used the natural log of the biomass, but untransformed biomass is presented here for ease of interpretation. N = 22 for no additions, 19 for algae only and algae supplement, 20 for mineral. Each rep is one data point

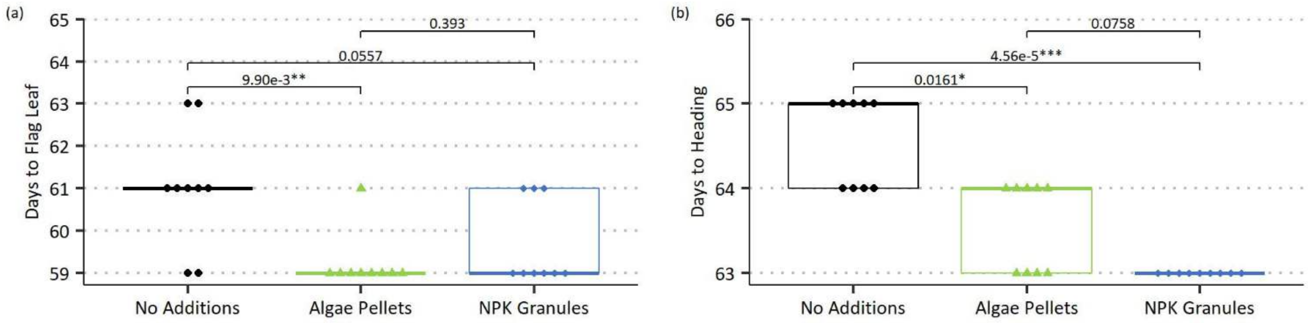

**Fig. S7 Key growth stages of cv. Laureate when grown in a field trial under three different fertiliser regimes. (a)** Days to half the plot with flag leaf unfurled. **(b)** Days to half the plot with ears emerged from the boot. For both boxplots, the thick horizontal line is the median, thin horizontal lines are the 1st and 3rd quartile. Overall significance determined by Kruskal-Wallis, p-value reported in the main text. Pairwise significance determined by Dunn test with Benjamini-Hochberg correction for multiple testing. N = 9 for all treatments. Each rep is one data point

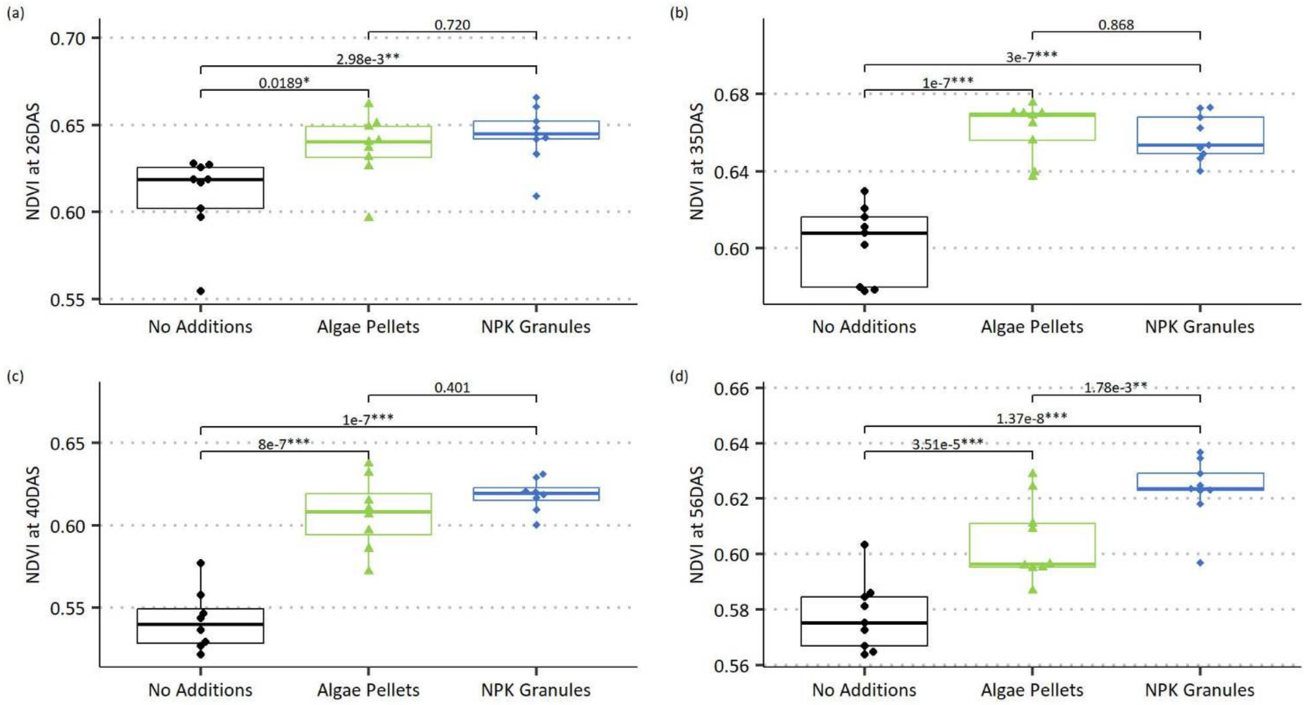

**Fig. S8 NDVI across the growth of cv. Laureate when grown in a field trial under three different fertiliser regimes. (a)** NDVI at 26DAS. **(b)** NDVI at 35DAS. **(c)** NDVI at 40DAS. **(d)** NDVI at 56DAS. For all boxplots, the thick horizontal line is the median, thin horizontal lines are the 1st and 3rd quartile. Whiskers extend to the range, excluding outliers (>1.5x the interquartile range below quartile 1 or above quartile 3). For **(a)**, **(b)** and **(c),** overall significance determined by ANOVA, p-value reported in the main text, and pairwise significance determined by Tukey's HSD. For **(d)**, the block effect was significant, so a mixed model was used. Overall significance determined by mixed model ANOVA, p-value reported in the main text, and pairwise significance determined by mixed model contrasts. N = 9 for all treatments. Each rep is one data point
